# Conditional lethality profiling reveals anticancer mechanisms of action and drug-nutrient interactions

**DOI:** 10.1101/2023.06.04.543621

**Authors:** Kyle M. Flickinger, Kelli M. Wilson, Nicholas J. Rossiter, Andrea L. Hunger, Tobie D. Lee, Matthew D. Hall, Jason R. Cantor

**Author notes:** Correspondence (J.R.C.). These authors contributed equally (Kyle M. Flickinger, Kelli M. Wilson). These authors contributed equally (Nicholas J. Rossiter, Andrea L. Hunger).

## Abstract

Chemical screening studies have identified drug sensitivities across hundreds of cancer cell lines but most putative therapeutics fail to translate. Discovery and development of drug candidates in models that more accurately reflect nutrient availability in human biofluids may help in addressing this major challenge. Here we performed high-throughput screens in conventional versus Human Plasma-Like Medium (HPLM). Sets of conditional anticancer compounds span phases of clinical development and include non-oncology drugs. Among these, we characterize a unique dual-mechanism of action for brivudine, an agent otherwise approved for antiviral treatment. Using an integrative approach, we find that brivudine affects two independent targets in folate metabolism. We also traced conditional phenotypes for several drugs to the availability of nucleotide salvage pathway substrates and verified others for compounds that seemingly elicit off-target anticancer effects. Our findings establish generalizable strategies for exploiting conditional lethality in HPLM to reveal therapeutic candidates and mechanisms of action.

## INTRODUCTION

Cell-based in vitro models are invaluable tools in drug discovery because they facilitate high-throughput screening based on defined molecular targets or phenotypes^1, 2^. Such models are also critical for preclinical drug development, as they enable studies that investigate mechanisms of action and evaluate how treatment responses vary with cell-intrinsic characteristics^3^. Despite the large increase in the catalogue of reported drug sensitivities across hundreds of human cancer cell lines, most experimental cancer therapeutics fail to translate – a problem attributed in part to the limited predictive value of preclinical models^4–8^. Thus, it is a central challenge to identify and interrogate drug candidates using models that more closely reflect conditions in the human body.

Traditional in vitro and in vivo models are useful for studying drug treatment phenotypes, but each has limitations. Most in vitro studies of human cell physiology and drug sensitivity have relied on growth media that poorly resemble the nutrient composition of human blood^9, 10^, incubators that expose cells to oxygen levels far greater than those in circulation and tissues^11, 12^, and culture formats that restrict most cell types to grow as 2D monolayers^13^. Direct in vivo screens for putative cancer drugs are impractical and cell growth conditions in animals are also difficult to control and manipulate, limiting opportunities to investigate environmental contributions to drug efficacy^14, 15^. There are also discrepancies in the nutrient composition of human versus mouse blood that could differentially affect cell metabolism and drug sensitivity^16^.

High-throughput phenotypic screening in cancer cells has enabled drug discovery without the need for prior knowledge of a specific target or mechanism of action^1, 2^. Efforts such as Genomics of Drug Sensitivity in Cancer (GDSC)^17, 18^, Cancer Target Discovery and Development (CTD2)^19, 20^, and DepMap^21^ have identified cancer therapeutic candidates using chemical screens across hundreds of cell lines, further revealing genomic determinants of drug sensitivity as well. Nonetheless, while recent work has shown that in vitro drug treatment phenotypes can depend on relative oxygen tension^22^, the presence of stromal cells^23, 24^, and culture in 3D models^25–28^, there has been considerably less attention on the influence of medium composition^29^ – arguably the most flexible component of the in vitro environment.

Previously, we developed Human Plasma-Like Medium (HPLM), a physiologic cell culture medium that contains over 60 components at concentrations that reflect their average values in adult human blood^16^. To create complete HPLM-based media, we add a 10% dialyzed fetal bovine serum (FBS) supplement (HPLM^+dS^) that contributes various growth factors, hormones, and trace elements required to broadly support cell growth, while minimizing the addition of undefined polar metabolites. Since RPMI 1640 (herein RPMI) is the historical medium of choice for culturing blood cells, we have also described RPMI-based reference media that each contain physiologic (5 mM) rather than RPMI-defined (11.1 mM) glucose and are further supplemented with either untreated (RPMI^+S^) or dialyzed (RPMI^+dS^) 10% FBS^16^. Guided by new findings in metabolic regulation and gene essentiality each linked to the availability of HPLM-specific metabolites, we have shown that culture in HPLM^+dS^ could alter responses to an approved chemotherapeutic agent (5-Fluorouracil; 5-FU) and to a compound (CB-839) that has been tested in cancer patients^16, 30^. Therefore, we reasoned that performing chemical screens in HPLM^+dS^ and RPMI-based media should facilitate the systematic identification of other compounds that differentially affect cell fitness under nutrient conditions with greater relevance to human physiology. By exploiting conditional lethality, it may also be possible to develop cancer treatment strategies based on coupling molecular therapeutics to the dietary or enzyme-mediated manipulation of circulating metabolites^31–33^.

Here, we perform high-throughput chemical screens to examine how medium composition affects drug sensitivity of human blood cancer lines. Analysis of these data reveal that conditional anticancer agents collectively span different phases of clinical development, vary with natural cell-intrinsic diversity, and include non-oncology drugs. Follow-up work traces conditional phenotypes for several chemotherapeutics to the availability of nucleotide salvage pathway substrates defined only in HPLM versus RPMI or otherwise contributed by the serum component of typical culture media. Additional analysis reveals a strikingly selective HPLM-sensitive phenotype for brivudine, a compound that has been approved for use as an antiviral agent in several European countries. By combining drug-nutrient complementation assays, metabolomics, CRISPR modifier screens, metabolite synthesis, and in vitro enzyme assays, we uncover a unique dual-mechanism model for conditional brivudine sensitivity. Using this highly integrative approach, we find that brivudine can disrupt de novo purine and thymidylate synthesis under low folate conditions by mediating inhibition of two cellular targets in folate metabolism. Moreover, we also leveraged our previous conditional gene essentiality data to reveal that canonical targets for other validated conditional anticancer compounds are encoded by nonessential genes. Together, these results demonstrate strategies for exploiting conditional lethality in HPLM to identify and examine cancer therapeutics and mechanisms of action with greater relevance to nutrient conditions in humans.

## RESULTS AND DISCUSSION

### High-throughput chemical screens reveal conditional anticancer compounds

Although chemical screens have identified drug candidates for a variety of diseases^1, 2, 4^, they have relied on growth media typically comprised of a synthetic basal component with limited relevance to nutrient levels reported in human blood and a serum supplement that contributes a cocktail of undefined biomolecules^9^. This is illustrated by cataloging the media used for phenotypic screens across hundreds of cell lines from the GDSC, CTD2, and DepMap projects. While these resources have uncovered many valuable insights, they are based on screens largely carried out in RPMI- or DMEM-based media supplemented with 5-20% FBS (Figure 1A)^17–21^. Given that we previously revealed two drug-nutrient interactions each linked to HPLM components not otherwise defined in traditional synthetic media, we reasoned that high-throughput screens could provide a powerful and unbiased approach for identifying other conditional anticancer compounds.

**Figure 1.**
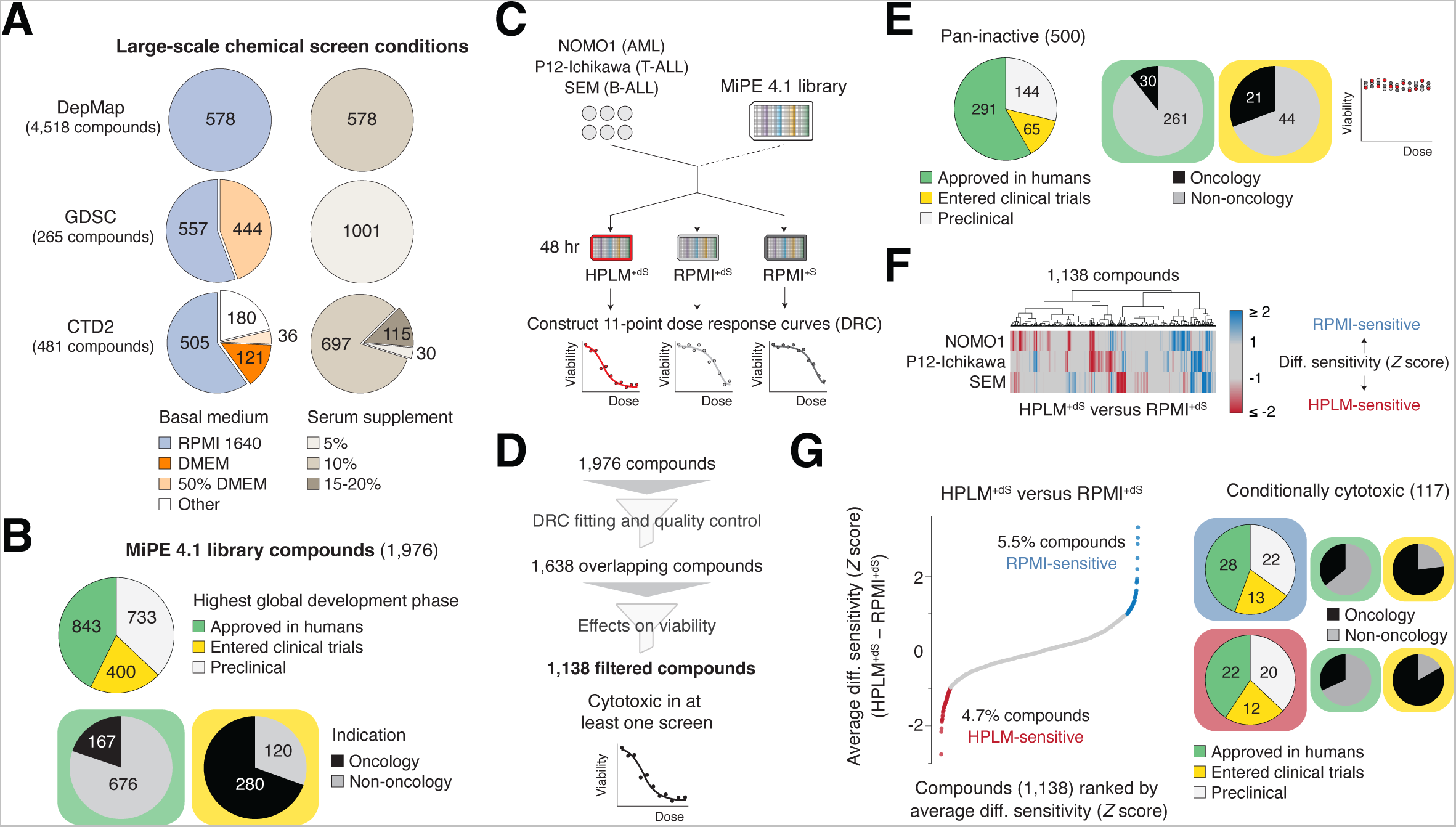
High-throughput screens for conditional anticancer compounds See also Figure S1; Table S1. (A) Growth conditions across chemical screens from the DepMap, Genomics of Drug Sensitivity in Cancer (GDSC), and Cancer Target Discovery and Development (CTD2) projects. 50% DMEM refers to basal media containing DMEM and another synthetic medium in a 1:1 mixture. (B) Highest global development phase and indication for compounds in the MIPE 4.1 library. (C) Schematic for high-throughput chemical screens in human blood cancer cells. AML, acute myeloid leukemia; ALL, acute lymphoblastic leukemia. (D) Schematic for dose-response curve quality control and filtering strategies to establish a set of overlapping compounds across screen datasets. See Methods. (E) Highest global development phase and indication for pan-inactive compounds. (F) Cluster map showing HPLM^+dS^ versus RPMI^+dS^ conditional phenotypes in three cell lines. (G) Compounds ranked by average HPLM^+dS^-RPMI^+dS^ sensitivity across three cell lines (left) (See methods). Highest global development phase and indication for conditionally sensitive hits (right).

To begin to pursue this, we used the NCATS Mechanism Interrogation Plate (MIPE) 4.1 library – a collection of 1,976 unique agents comprised of investigational compounds and drugs that have either entered clinical testing or been approved for use to treat cancer and a variety of non-oncology indications (Figures 1B and S1A; Table S1)^34^. We screened a panel of three blood cancer cell lines (NOMO1, acute myeloid leukemia (AML); P12-Ichikawa, T-cell acute lymphoblastic leukemia (ALL); SEM, B-cell ALL) against each MIPE compound over an eleven-point concentration range by measuring viability following 48 hr treatment in HPLM^+dS^, RPMI^+dS^, or RPMI^+S^ (Figure 1C). We then systematically generated dose-response curves and applied filtering strategies to minimize possible artifacts and to, in turn, remove a set of 500 pan-inactive compounds that did not elicit cytotoxic effects in any of the screens (Figures 1D, 1E, and S1B).

For each of the 1,138 filtered compounds shared across all screen datasets, we defined the response score as the area under the dose-response curve (AUC). Interestingly, these scores were more strongly correlated by cell line than screen condition (Figure S1C). To establish conditional sensitivity profiles, we then standardized (Z-score) each set of differential response scores: (1) HPLM^+dS^-RPMI^+dS^; (2) HPLM^+dS^-RPMI^+S^; and (3) RPMI^+dS^-RPMI^+S^. Overall, these profiles showed variable degrees of overlap between cell lines (Figures 1F and S1D). Given the size of our panel, we then averaged each respective profile across cell lines to increase detection of the strongest hits. We first asked how the differential availability of defined medium components affected drug treatment responses. By setting a Z-score cutoff of 1, our HPLM^+dS^-RPMI^+dS^ profile revealed 54 HPLM-sensitive and 63 RPMI-sensitive compounds (Figure 1G). Only one-third of these hits have been approved or tested for cancer treatment in humans, while the remainder were comprised of investigational compounds and non-oncology drugs.

### Drug-nutrient interactions between purine analogs and hypoxanthine

The top-three scoring RPMI-sensitive hits from this profile were purine analogs that have each been approved for cancer therapy – dacarbazine (DTIC), 6-Mercaptopurine (6-MP), and 6-Thioguanine (6-TG) (Figure 2A). Upon cell entry, 6-MP and 6-TG are enzymatically converted to active metabolites that inhibit de novo purine synthesis or become misincorporated into nucleic acids, while DTIC is often described as a DNA alkylating agent that is first metabolically activated in the liver (Figure S2A)^35–37^. These conditional phenotypes were not conserved in the HPLM^+dS^-RPMI^+S^ profile, suggesting that they may be linked to a nutrient(s) whose defined differential availability could be complemented by the FBS (Figure 2B). 6-MP and 6-TG are non-endogenous substrates for HPRT, which converts both guanine to GMP and hypoxanthine to IMP in the purine salvage pathway^38^. Guanine is not a defined component in either synthetic medium, but HPLM contains hypoxanthine at a concentration similar to levels provided by 10% FBS prior to dialysis (Figure 2C). Therefore, we hypothesized that conditional sensitivities to 6-MP and 6-TG might be traced to hypoxanthine, which could compete with both drugs as a substrate for HPRT.

**Figure 2.**
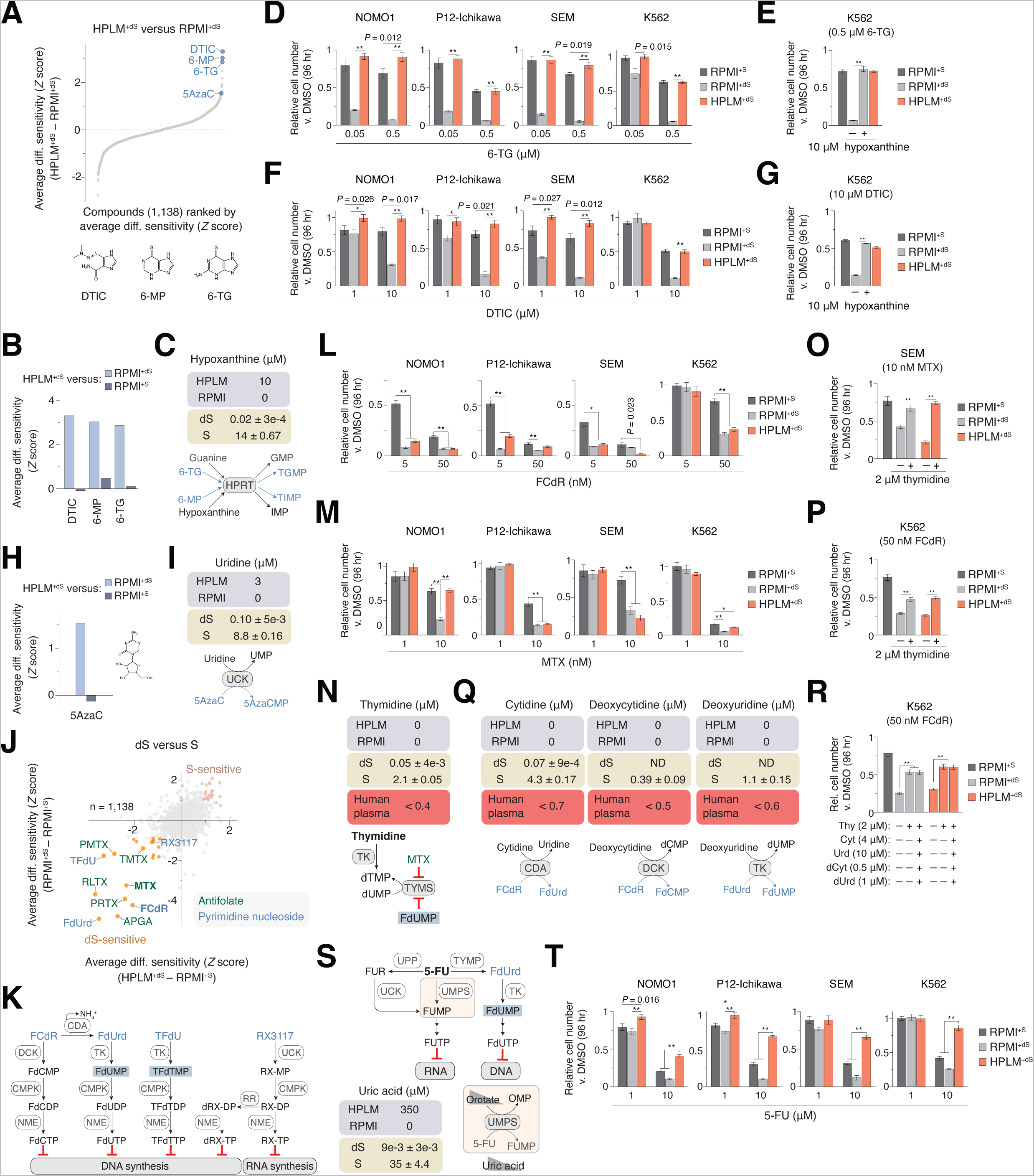
Drug-nutrient interactions with nucleotide salvage pathway substrates See also Figure S2; Tables S1 and S3. (A) Compounds ranked by average HPLM^+dS^-RPMI^+dS^ sensitivity. DTIC, dacarbazine; 6-MP, 6-Mercaptopurine; 6-TG, 6-Thioguanine; 5AzaC, 5-Azacytidine. (B) Conditional phenotypes for DTIC, 6-MP, and 6-TG from averaged HPLM^+dS^-RPMI^+dS^ and HPLM^+dS^-RPMI^+S^ profiles. (C) Defined hypoxanthine levels in HPLM and RPMI. Concentrations of hypoxanthine in 10% FBS (dS, dialyzed; S, untreated) (mean ± SD, *n* = 3) (top). Schematic of reactions catalyzed by HPRT, hypoxanthine-guanine phosphoribosyltransferase (bottom). (D-G) Relative growth of cells treated with 6-TG (D and E) or DTIC (F and G) versus DMSO (mean ± SD, *n* = 3, \*\**P* < 0.005, * *P* < 0.01). (H) Conditional phenotypes for 5AzaC from averaged HPLM^+dS^-RPMI^+dS^ and HPLM^+dS^-RPMI^+S^ profiles. (I) Defined uridine levels in HPLM and RPMI. Concentrations of uridine in 10% FBS (dS, dialyzed; S, untreated) (mean ± SD, *n* = 3) (top). Schematic of reactions catalyzed by UCK, uridine-cytidine kinase (bottom). (J) Comparison of averaged HPLM^+dS^-RPMI^+S^ and RPMI^+dS^-RPMI^+S^ phenotypes across the three screened cell lines. dS-sensitive pyrimidine nucleoside analogs (blue) and antifolates (green) are labeled. FCdR, fluorodeoxycytidine; MTX, methotrexate. Remaining compound abbreviations in Figure S2. (K) Cellular conversion of dS-sensitive pyrimidine nucleoside analogs to metabolites that mediate cytotoxic effects. Fluorodeoxyuridine monophosphate (FdUMP) and trifluoromethyl deoxyuridine monophosphate (TFdTMP) inhibit thymidylate synthase (TYMS). (L-M) Relative growth of cells treated with FCdR (L) or MTX (M) versus DMSO (mean ± SD, *n* = 3, \*\**P* < 0.005). (N) Thymidine is not defined in HPLM or RPMI. Concentrations of thymidine in 10% FBS (dS, dialyzed; S, untreated) (mean ± SD, *n* = 3) (top). Thymidine availability reported for human plasma (middle). Reactions catalyzed by thymidine kinase (TK) and TYMS that generate dTMP (bottom). dUMP, deoxyuridine monophosphate; dTMP, deoxythymidine monophosphate. (O-P) Relative growth of cells treated with MTX (O) or FCdR (P) versus DMSO (mean ± SD, *n* = 3, \*\**P* < 0.005). (Q) Cytidine, deoxycytidine, and deoxyuridine are not defined in HPLM or RPMI. Concentrations of each in 10% FBS (dS, dialyzed; S, untreated) (mean ± SD, *n* = 3) (top) and levels reported for human plasma (middle)^47, 125^. Reactions catalyzed by enzymes that can act on these pyrimidines and various FCdR derivatives (bottom). (R) Relative growth of cells treated with FCdR versus DMSO (mean ± SD, *n* = 3, \*\**P* < 0.005). (S) Schematic of 5-fluorouracil (5-FU) metabolism (top). Distinct branch-points generate FdUMP (and FdUTP) or FUTP, with the latter initiated by UMPS. Uric acid is an endogenous inhibitor of UMPS and its relative availability can affect cellular levels of orotate, which competes with 5-FU as a substrate for UMPS^16^ (bottom). (T) Relative growth of cells treated with 5-FU versus DMSO (mean ± SD, *n* = 3, \*\**P* < 0.005, \**P* < 0.01).

To interrogate conditional phenotypes, we measured cell counts from suspension cultures following treatment with each of two doses over a ten-fold concentration range selected to capture possible medium-dependent responses. Given the potential to integrate data from our recent work with CRISPR-based conditional gene essentiality profiling^30^, we extended these follow-up assays to the K562 chronic myeloid leukemia (CML) cell line – further testing dose-responses in this line over extended six-point concentration ranges.

We first validated the conditional phenotype for 6-TG in all four cell lines, as at least one tested dose elicited 60-80% growth defects in RPMI^+dS^ versus both HPLM^+dS^ and RPMI^+S^ (Figures 2D and S2B). When we supplemented RPMI^+dS^ with HPLM-defined hypoxanthine levels (10 μM), we observed complete rescue of these defects in K562 cells as anticipated (Figure 2E). We then confirmed that conditional phenotypes for DTIC across cell lines were similar to those for 6-TG but required higher treatment doses, suggesting that DTIC can also act as a weaker purine analog competing with hypoxanthine or perhaps a de novo purine synthesis substrate beyond its activity as an alkylating agent (Figures 2F and S2C). Equivalent addition of physiologic hypoxanthine to RPMI^+dS^ indeed normalized K562 cell sensitivities to DTIC across conditions (Figure 2G). Notably, our previous genome-wide CRISPR screens in HPLM^+dS^ versus RPMI^+dS^ revealed that four of the six genes encoding enzymes along the de novo purine synthesis pathway were RPMI-essential in K562 cells – phenotypes also likely linked to hypoxanthine availability (Figure S2D). Together, these results suggest that targeting systemic hypoxanthine availability may provide opportunities to increase the potency of purine analogs and putative inhibitors of de novo purine synthesis.

Since uridine is another component specific to HPLM versus RPMI, we then considered if any pyrimidine analogs might be RPMI-sensitive hits as well. 5-Azacytidine (5AzaC) is approved to treat certain cancers and was indeed also among the top 20% of positive hits from our HPLM^+dS^-RPMI^+dS^ profile. Similar to the purine analogs above, this conditional phenotype for 5AzaC was not seen in the HPLM^+dS^-RPMI^+S^ profile (Figure 2H). 5AzaC is converted by cellular enzymes to effector metabolites that disrupt nucleic acid synthesis (Figure S2E)^36^. This activation is initiated by UCK, which also catalyzes the conversion of uridine to UMP in the pyrimidine salvage pathway. Similar to the case for hypoxanthine, uridine levels provided in 10% FBS are 100-fold greater prior to dialysis – equivalent to a concentration 3-fold greater than that defined in HPLM (3 μM) (Figure 2I). Therefore, these conditional phenotypes can likely be traced to uridine, which can antagonize the conversion of 5AzaC to active metabolites.

### Serum thymidine affects cell sensitivity to TYMS inhibitors

Next, we explored whether FBS might further influence drug sensitivity regardless of basal medium. By setting a Z-score cutoff of 0.9, we identified 19 dS-sensitive and 20 S-sensitive hits based on shared conditional phenotypes over the HPLM^+dS^-RPMI^+S^ and RPMI^+dS^-RPMI^+S^ profiles (Figure 2J). Interestingly, most S-sensitive compounds were investigational, while more than half of the dS-sensitive hits were comprised of antifolates and pyrimidine nucleoside analogs that have largely been approved or tested as chemotherapeutics (Figure S2F). Antifolates can either directly or indirectly inhibit enzymes involved in folate metabolism, including thymidylate synthase (TYMS) and dihydrofolate reductase (DHFR), while the analog hits are converted to fluoronucleotides that either interfere with nucleic acid synthesis or inhibit TYMS (Figures 2K and S2G)^39–43^. We sought to investigate conditional phenotypes for one compound from each of these dS-sensitive subsets. Given its broad set of effector metabolites, we first examined fluorodeoxycytidine (FCdR).

For at least one tested dose, we confirmed that FCdR induced much stronger growth defects in the dS-supplemented media versus RPMI^+S^ across all four cell lines (Figures 2L and S2H). Among the antifolates, we selected methotrexate (MTX) because it remains widely used for the treatment of various cancers worldwide^38, 44^. However, conditional phenotypes for MTX varied between cell lines, with 30-50% impaired growth in either both dS-containing media (SEM, P12-Ichikawa) or in RPMI^+dS^ alone (NOMO1), or instead, more modest dS-dependent responses that were greater in RPMI^+dS^ relative to HPLM^+dS^ (K562) (Figures 2M and S2I).

Since TYMS inhibition is a mechanism shared among several antifolates and pyrimidine nucleoside analogs, we considered if this activity might contribute to the dS-sensitive phenotypes. Notably, our previous secondary CRISPR screens in the K562 line revealed that *TYMS* was a strong HPLM-essential hit versus RPMI^+S^ but not RPMI^+dS^, reflective of a comparable dS-essential phenotype (Figure S2J). TYMS catalyzes the conversion of 5,10-methylene-tetrahydrofolate (5,10-meTHF) and dUMP to dihydrofolate (DHF) and thymidylate (dTMP). This reaction serves as the only de novo source of dTMP, which can be further metabolized to dTTP – an essential substrate in DNA synthesis and repair pathways^40^. Similar to the nucleotide salvage reactions that generate IMP and UMP, there are two human thymidine kinases that can convert thymidine to dTMP but differ in subcellular localization (TK1, cytosolic; TK2, mitochondrial)^45^. Thymidine is not defined in traditional synthetic media and also failed to meet inclusion criteria set in our design of HPLM on the basis of its sub-micromolar availability in human blood^46, 47^. However, 10% FBS provides supraphysiologic thymidine levels (2 μM) that are reduced by more than 40-fold following dialysis (Figure 2N). Therefore, we hypothesized that conditional phenotypes for MTX and FCdR may be linked to FBS-contributed thymidine, which can serve as a salvage source of dTMP when TYMS activity is compromised.

To test this, we added 2 μM thymidine to RPMI^+dS^ or HPLM^+dS^, and indeed found that SEM cell responses to MTX were normalized versus RPMI^+S^ (Figure 2O). These results are consistent with prior work demonstrating that thymidine availability could affect MTX toxicity in cultured cells, mouse models, and patients^48–50^. Of note, MTX is also a potent DHFR inhibitor. Consistent with the notion that cellular folates further support de novo purine biosynthesis, evidence has shown that impaired growth from either chemical or genetic inhibition of DHFR may be rescued using a combined supplementation of thymidine and hypoxanthine^51–54^. Thus, we reason that conditional MTX treatment phenotypes can be influenced by an interplay of differential demands for purines and thymidylate, cell-intrinsic utilization of nucleotide salvage pathways, and protective effects provided by HPLM-defined hypoxanthine.

In contrast to our results for MTX, equivalent thymidine supplementation provided only a 50% rescue of relative growth defects in FCdR-treated K562 cells (Figure 2P). Among the effector metabolites of FCdR and a few other pyrimidine analogs is fluorodeoxyuridine monophosphate (FdUMP), which forms a covalent ternary complex with TYMS and 5,10-meTHF^55–57^. Despite the incomplete rescue in thymidine-supplemented conditions, FdUMP was the only fluoronucleotide that we could detect in FCdR-treated K562 cells by liquid chromatography-mass spectrometry (LC-MS) (Figure S2K). Therefore, we considered the possibility that other FCdR derivatives were below MS detection limits or incorporated into nucleic acids, and in turn, if FBS contributes other pyrimidines that may affect FCdR activation. While we detected three additional pyrimidines that could antagonize different steps in FCdR metabolism (cytidine, deoxycytidine, and deoxyuridine), their combined supplementation to the dS-containing media offered no further rescue, suggesting that FCdR sensitivity can be influenced by additional but unanticipated FBS components as well (Figures 2Q and 2R).

5-FU is another pyrimidine base analog used as a cancer therapeutic^38^. One branch-point of 5-FU activation leads to the formation of FdUMP from floxuridine (FdUrd)^58^ – a precursor shared with FCdR metabolism and also a compound identified among the set of dS-sensitive hits (Figure 2S). Indeed, the primary mechanism of action for 5-FU is often attributed to TYMS inhibition^40, 41^. However, we previously showed that 5-FU sensitivity in NOMO1 cells was reduced in HPLM^+dS^ versus both RPMI-based media, effects traced to uric acid availability and a distinct metabolic branch-point of 5-FU activation^16^. Notably, this RPMI-sensitive phenotype was further shared across our cell line panel (Figures 2T and S2L). Together, these results suggest that 5-FU sensitivity may depend on an interplay between uric acid availability and the cell-dependent expression of TYMP, the enzyme that converts 5-FU to FdUrd.

### Identification of a drug-nutrient interaction between brivudine and folic acid

After excluding dS- and S-sensitive hits from further analysis, we used our data to uncover conditional anticancer compounds regardless of the RPMI-based reference medium. By setting a Z-score cutoff of 0.9, we identified 26 RPMI-sensitive and 26 HPLM-sensitive compounds based on shared conditional phenotypes in the HPLM^+dS^-RPMI^+dS^ and HPLM^+dS^-RPMI^+S^ profiles (Figure 3A). Among these, the top-scoring hit was brivudine (BVDU), an antiviral agent against herpes simplex virus type 1 (HSV-1) and varicella-zoster virus (VSV) approved to treat herpes zoster in several European countries^59, 60^. Our screen results suggested that BVDU was HPLM-sensitive and not differentially toxic between the RPMI-based media, suggesting that undefined metabolites in FBS had little impact on relative BVDU sensitivity (Figure S3A). Upon entry into virally infected cells, it has been reported that BVDU can be converted to effector metabolites that interfere with viral DNA synthesis or inhibit TYMS^60–62^. Canonical BVDU selectivity against virally infected cells is attributed to the requirement for HSV/VSV-encoded TK to catalyze reactions that convert BVDU to its mono-(BVDU-MP) and diphosphate (BVDU-DP) forms^61, 63, 64^. BVDU-DP is further converted to the active triphosphate metabolite (BVDU-TP) that disrupts viral DNA synthesis, while it has also been suggested that BVDU-MP can inhibit the viral TYMS (Figure 3B)^61, 65^. Notably, the two human TKs show marked differences in kinase activity for various natural and non-endogenous substrates, including a TK2-specific affinity for BVDU, thus suggesting a potential non-viral source of BVDU-MP in BVDU-treated human cells^66–68^.

**Figure 3.**
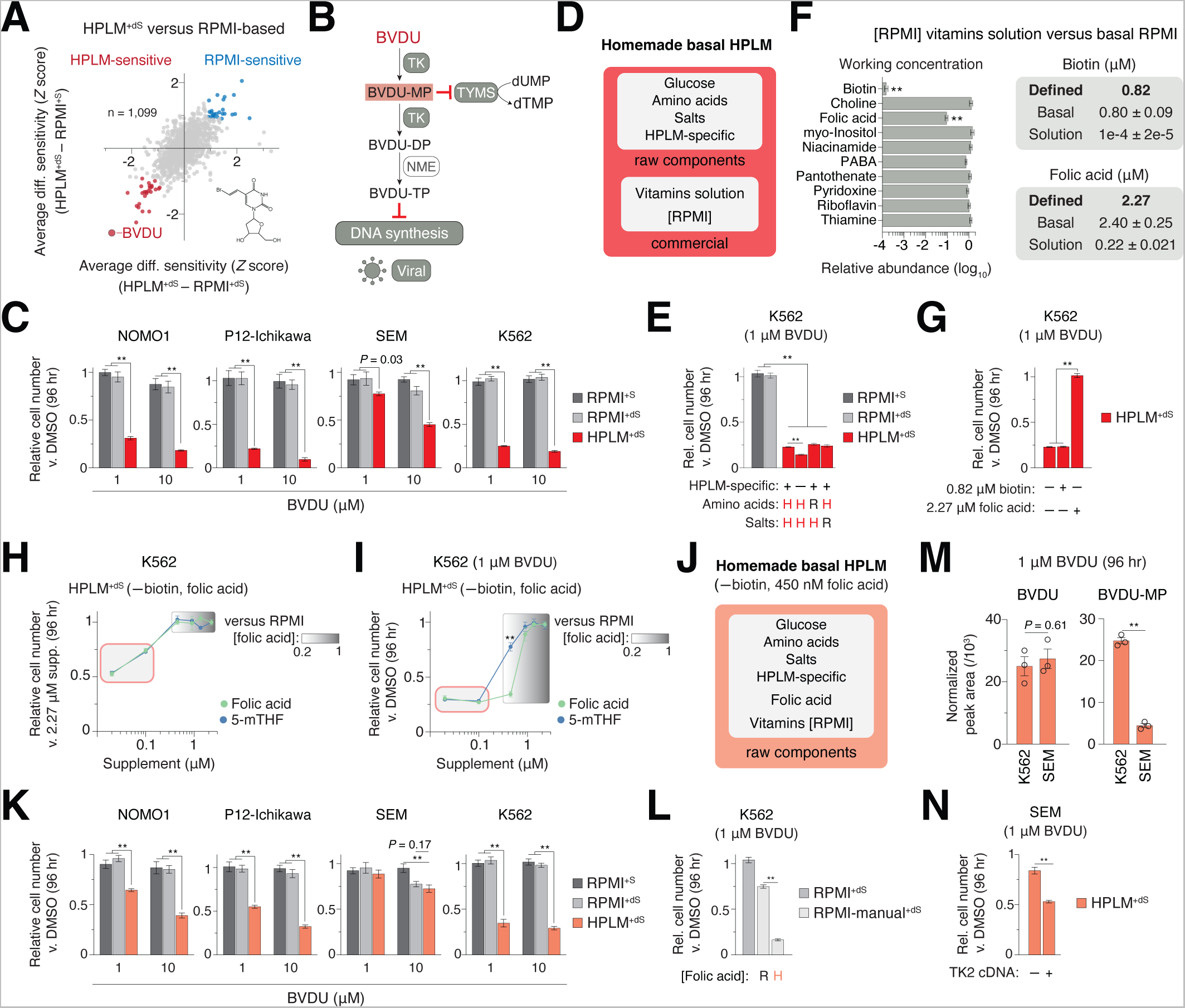
Identification of a drug-nutrient interaction between brivudine and folic acid See also Figure S3; Tables S1, S2, and S3. (A) Comparison between averaged HPLM^+dS^-RPMI^+dS^ and HPLM^+dS^-RPMI^+S^ phenotypes across the three screened cell lines. Brivudine (BVDU) is labeled. (B) Schematic for the activation and canonical mechanism of BVDU action in virally infected cells. Viral TK catalyzes reactions that convert BVDU to its mono-(BVDU-MP) and diphosphate (BVDU-DP) forms. BVDU-MP can inhibit viral TYMS. BVDU-DP is metabolized to the active triphosphate derivative (BVDU-TP) that inhibits viral DNA polymerase. (C) Relative growth of cells treated with BVDU versus DMSO (mean ± SD, *n* = 3, \*\**P* < 0.005). (D) Components of homemade HPLM. (E) Relative growth of cells treated with BVDU versus DMSO (mean ± SD, *n* = 3, \*\**P* < 0.005). H, HPLM-defined concentrations. R, RPMI-defined concentrations. See Table S2 for further detail regarding construction of RPMI-defined component stocks. (F) Relative working concentrations of vitamins in RPMI 100X vitamin solution (Sigma R7256, Lot RNBB7627) versus basal RPMI (Thermo 11879, Lot 2458379) (mean ± SD, *n* = 3, \*\**P* < 0.005) (left). Defined and working concentrations of biotin and folic acid (mean ± SD, *n* = 3) (right). RPMI also contains 3.69 nM vitamin B12, which could not be detected by the profiling method. (G) Relative growth of cells treated with BVDU versus DMSO (mean ± SD, *n* = 3, \*\**P* < 0.005). (H) Relative growth of K562 cells in HPLM^+dS^ (lacking biotin and folic acid) with increasing levels of folic acid or 5-mTHF versus the respective supplement at a concentration equal to that of RPMI-defined folic acid (2.27 μM) (mean ± SD, *n* = 3). Red shaded box, range reported for these folates in human plasma (See Figure S3). Gradient shaded box, range from 450 nM to 2.27 μM. 5-mTHF, 5-methyltetrahydrofolate. (I) Relative growth of K562 cells treated with BVDU versus DMSO in HPLM^+dS^ (lacking biotin and folic acid) with increasing levels of folic acid or 5-mTHF (mean ± SD, *n* = 3, \*\**P* < 0.005). (J) Components of a modified homemade HPLM that lacks biotin and contains 450 nM folic acid. HPLM-based media are distinguished with the shaded colors in (D) and (J). Of note, this modified basal HPLM was used for all experiments in this study except the chemical screens and those in panels (C), (E), (G) and (S3B). (K) Relative growth of cells treated with BVDU versus DMSO (mean ± SD, *n* = 3, \*\**P* < 0.005). (L) Relative growth of cells treated with BVDU versus DMSO (mean ± SD, *n* = 3, \*\**P* < 0.005). H, HPLM-defined concentration (450 nM). R, RPMI-defined concentration (2.27 μM). See Table S2 for further detail regarding construction of homemade (manual) RPMI. (M) Cellular abundances of BVDU and BVDU-MP in the K562 and SEM lines following treatment with BVDU in HPLM^+dS^ (mean ± SD, *n* = 3, \*\**P* < 0.005). (N) Relative growth of cells treated with BVDU versus DMSO (mean ± SD, *n* = 3, \*\**P* < 0.005).

BVDU elicited strikingly selective 70-90% growth defects specific to HPLM^+dS^ in three cell lines, and more modest (30-40%) HPLM-dependent cytotoxicity in SEM cells (Figure 3C). Further, when we tested BVDU against K562 cells across an eight-point concentration range, we found that even the lowest dose (10 nM) impaired growth by 30% in HPLM^+dS^, while the highest one (50 μM) caused only minor growth defects (< 10%) in RPMI-based media (Figure S3B). To determine why cells were selectively sensitive to BVDU in HPLM^+dS^, we initially considered links to effector metabolites involved in antiviral BVDU activity. Since the VSV and human TYMS homologs share over 80% sequence homology, we reasoned that TK2 could perhaps facilitate TYMS inhibition by catalyzing BVDU-MP synthesis. However, this mechanism alone was not consistent with the lack of BVDU cytotoxicity observed in RPMI^+dS^, which contains negligible thymidine similar to HPLM^+dS^. Moreover, conditional phenotypes for BVDU were distinct from those for TYMS inhibitors among the set of dS-sensitive hits. Therefore, the underlying cause for conditional BVDU cytotoxicity was not immediately apparent.

Basal HPLM contains glucose, proteinogenic amino acids, salts, and over thirty additional components not otherwise defined in traditional synthetic media (Table S2). Additionally, several vitamins are essential for mammalian cell culture but failed to meet inclusion criteria set in our design of HPLM^16, 69^. Rather than omit these important nutrients, we had used a commercial stock solution (Sigma R7256) to incorporate vitamins into HPLM at RPMI-defined levels (Figure 3D). Thus, given the availability of glucose and vitamins were presumed to be normalized between the two basal media, we systematically tested BVDU against K562 cells in HPLM-based derivatives that contained remaining basal components adjusted to reflect their RPMI-defined concentrations. Each of these derivatives failed to rescue BVDU-induced toxicity, and interestingly, growth defects were modestly exacerbated when the HPLM-specific metabolites were removed (Figure 3E).

Although we expected vitamin concentrations to be equivalent in homemade HPLM versus RPMI, we previously uncovered discrepancies through metabolite profiling of complete media^16, 30^. Since we prepare the glucose added to our HPLM- and RPMI-based media, we then hypothesized that conditional BVDU sensitivity could be linked to unforeseen differences in vitamin availability. To test this, we first profiled vitamins in basal RPMI and the commercial stock solution and then determined their respective concentrations in complete media. While working levels for most were comparable between the two reagents, those for biotin and folic acid were 10,000- and 10-fold lower, respectively, only as provided by the stock solution relative to defined values (Figure 3F). Indeed, when we created HPLM derivatives with RPMI-defined levels of these vitamins, we found that folic acid completely rescued the growth defect of BVDU-treated K562 cells (Figure 3G).

Folic acid and 5-methyl-THF (5-mTHF) are the two most abundant folates in human blood, but circulating levels of each are up 100-fold lower than the RPMI-defined folic acid concentration (Figure S3C)^70–73^. Therefore, we next examined how the relative availability of these folates affect BVDU sensitivity. We created biotin- and folic acid-free HPLM that contained the eight remaining MS-detectable RPMI vitamins from manually prepared stocks, and then systematically added folic acid or 5-mTHF at concentrations spanning either reported values in human plasma (20-100 nM) or a five-fold range relative to RPMI-defined folic acid (0.45-2.27 μM). Across these derivatives, the two folates had nearly identical effects on the relative growth of K562 cells, with comparable 25-50% defects seen only at the plasma-like concentrations (Figure 3H). In contrast, while BVDU impaired cell growth across all derivatives containing physiologic folate levels, 5-mTHF began to protect against these defects at a concentration (450 nM) 2-fold lower than was required for folic acid – perhaps given that SLC19A1, the major folate importer in mammalian cells^74^, has a much stronger affinity for 5-mTHF (Figure 3I)^75^.

To preserve the conditional phenotype for BVDU while minimizing baseline growth defects from plasma-like folate availability, we used our modified HPLM with 450 nM folic acid for follow-up work (Figure 3J). When we evaluated BVDU sensitivity in this HPLM derivative, we found that growth defects across cell lines were partially reduced versus those seen in the original HPLM^+dS^, as could perhaps be expected given the 2-fold boost in folic acid provided (Figures 3K and S3D). In addition, reducing folic acid to 450 nM in RPMI created with manually prepared components was sufficient to dramatically increase K562 cell responses to BVDU in RPMI^+dS^ (Figure 3L).

Since SEM cells showed a much lower relative BVDU sensitivity regardless of the basal HPLM tested, we hypothesized that uptake or putative activation of BVDU in these cells may be reduced relative to the other cell lines. To test this, we sought to measure the levels of free and phosphorylated forms of BVDU in SEM and K562 cells following treatment in HPLM^+dS^. Whereas the BVDU levels were comparable, those for BVDU-MP were 5-fold greater in K562 cells, while BVDU-DP and -TP could not be detected in any samples (Figure 3M). Whether or not the latter two metabolites were below MS detection limits, these results suggested that BVDU anticancer activity depends on metabolic synthesis of BVDU-MP as perhaps mediated by TK2 in cells lacking viral TK. Consistent with this, stable expression of a *TK2* cDNA increased the growth defects of BVDU-treated SEM cells by 30% (Figures 3N and S3E).

### Brivudine interferes with folate-dependent nucleotide synthesis

Folate metabolism serves to activate and transfer one-carbon (1C) units that support many biochemical processes, including de novo purine and thymidylate synthesis, mitochondrial protein translation, and methionine regeneration^76, 77^. Within this broad network of 1C metabolism, folates function as the carriers of attached 1C units (e.g., methyl and formyl groups). In mammalian cells, there are parallel 1C pathways in the cytosol and mitochondria that are typically interlinked such that the mitochondrial 1C pathway produces formate, which is exported to the cytosol and used to support the synthesis of nucleotides and methionine^76, 77^.

Since our HPLM derivative constructed with RPMI-defined levels of methionine and other amino acids had limited impact on BVDU sensitivity, we considered the possibility that conditional BVDU cytotoxicity might be linked to folate-dependent nucleotide synthesis. Nonetheless, given our earlier rationale that TYMS was unlikely the primary target of BVDU, we first sought to confirm that thymidine supplementation is sufficient to rescue impaired growth specific to TYMS inhibition. Therefore, we engineered *TYMS*-knockout K562 cells, which showed growth defects in HPLM^+dS^ that were indeed completely rescued by the addition of 4 μM thymidine but unaffected by a 4-fold increase in hypoxanthine availability (40 μM) (Figures 4A and S4A). In contrast, supplementation with up to 8 μM thymidine had little impact on BVDU sensitivity in K562 cells, while hypoxanthine up to 40 μM provided dose-dependent but only partial protective effects (Figures 4B and S4B). However, the combined addition of supraphysiologic hypoxanthine (40 μM) and thymidine (4 μM) to HPLM^+dS^ completely rescued BVDU-induced cytotoxicity. Together, these results suggest that the BVDU-folic acid interaction can be traced to the hierarchical inhibition of de novo purine and dTMP synthesis, likely explaining why BVDU toxicity was exacerbated in our derivative lacking hypoxanthine and other HPLM-specific components. Consistent with this notion, BVDU markedly reduced the cellular levels of ATP, GTP, and dTTP but not of CTP and UTP – changes that were largely reversed in HPLM^+dS^ containing RPMI-defined folic acid (Figures 4C and S4C). Notably, this 5-fold boost in folic acid had little effect on BVDU-MP levels, suggesting that the drug-nutrient interaction was not linked to relative BVDU activation (Figure S4D). Moreover, K562 cells did not show any net folic acid exchange regardless of BVDU treatment or initial folic acid availability in HPLM, suggesting that this interaction could not be attributed to a limiting depletion of exogenous folic acid as well (Figure S4E).

**Figure 4.**
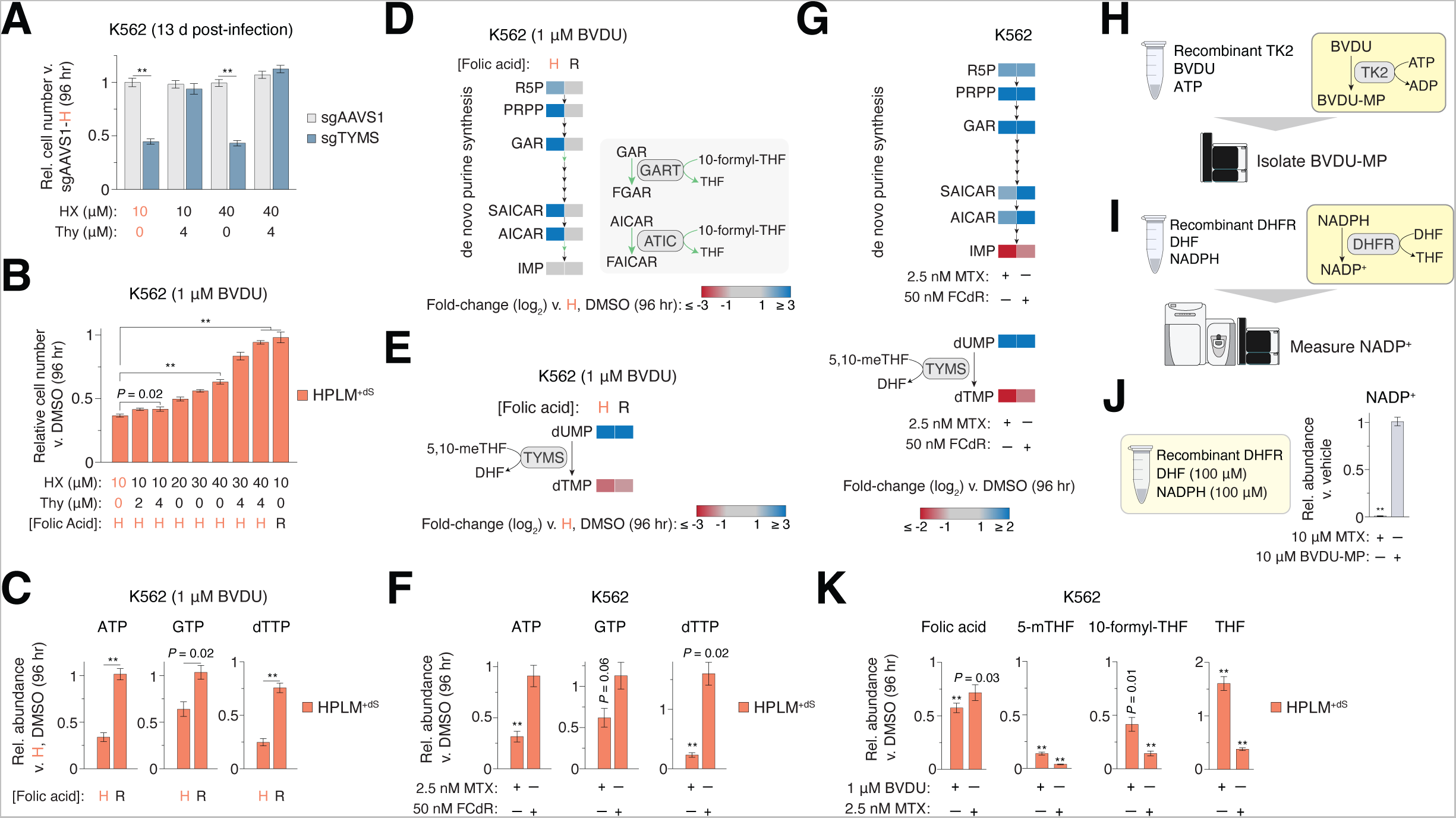
Brivudine interferes with folate-dependent nucleotide synthesis See also Figure S4; Tables S3. (A) Relative growth of *TYMS*-knockout and control cells versus control cells in HPLM^+dS^ (mean ±SD, *n* = 3, \*\**P* < 0.005). (B) Relative growth of cells treated with BVDU versus DMSO (mean ± SD, *n* = 3, \*\**P* < 0.005). H, HPLM-defined concentration. R, RPMI-defined concentration. (C) Relative abundances of ATP, GTP and dTTP in BVDU-treated and control cells versus control cells in HPLM^+dS^ (mean ± SD, *n* = 3, \*\**P* < 0.005). (D) Heatmap of relative abundances for metabolites along the de novo purine synthesis pathway in BVDU-treated and control cells versus control cells in HPLM^+dS^ (left). Two steps in this pathway use 10-formyl-THF as a co-substrate (right). (E) Heatmap of relative abundances for two TYMS reaction components (left). BVDU-treated and control cells versus control cells in HPLM^+dS^. (F) Relative abundances of ATP, GTP and dTTP in FCdR- and MTX-treated versus control cells in HPLM^+dS^ (mean ± SD, *n* = 3, \*\**P* < 0.005). Concentrations for FCdR and MTX were selected to elicit growth defects comparable to those for BVDU-treated K562 cells in HPLM^+dS^. (G) Heatmap of relative abundances for metabolites highlighted in (D) and (E). FCdR- and MTX- treated versus control cells in HPLM^+dS^. (H) Schematic for a method to isolate BVDU-MP from reactions catalyzed by human TK2. (I) Schematic of an assay for DHFR activity using LC-MS. (J) Relative NADP abundances versus vehicle for the assay in (I) (mean ± SD, *n* = 3, \*\**P* < 0.005). (K) Relative abundances of folate metabolites in BVDU- and MTX-treated versus control cells in HPLM^+dS^ (mean ± SD, *n* = 3, \*\**P* < 0.005).

Unbiased metabolite profiling revealed that BVDU had additional effects on the K562 cell metabolome (Figure S4F; Table S3). Among the most prominent were 10- to 20-fold increases in the abundances of several de novo purine synthesis intermediates, which may be attributed to a deficiency in cytosolic 10-formyl-THF – the folate species that feeds into this pathway at each of two steps (Figure 4D)^78^. Of note, these changes were coupled to a more modest 40% reduction in IMP (Table S3). In addition, BVDU treatment caused a 90-fold increase in dUMP levels and a 5-fold reduction in dTMP – the pyrimidine components of the TYMS reaction (Figure 4E). Overall, BVDU-induced changes to the metabolome were largely reversed by the addition of RPMI-defined folic acid, though dUMP abundances were still 15-fold greater relative to untreated cells – effects parallel to an incomplete normalization of dTMP and dTTP levels as well. These results suggested that increased folic acid availability relieves differential impairment thresholds on the two folate-dependent nucleotide synthesis pathways in BVDU-treated cells.

FCdR shares structural similarity with BVDU and MTX effectively inhibits de novo purine and thymidylate synthesis as well. Thus, we next asked how these drugs affect the same subsets of metabolites following treatment of K562 cells with doses that elicit comparable growth defects in HPLM^+dS^ (Figure S4G). Interestingly, FCdR had little impact on ATP and GTP levels but induced 60-90% increases in the abundances of dTTP, CTP, and UTP – effects that could be attributed to disrupted incorporation of pyrimidines into nucleic acids – while changes across this NTP panel in MTX-treated cells resembled those seen for BVDU (Figures 4F and S4H). Nonetheless, relative abundance effects across de novo purine intermediates and TYMS reaction components in both cases largely phenocopied those seen in BVDU-treated cells (Figure 4G). Together, these results were consistent with the incomplete rescue of FCdR-treated cells in dS-supplemented media with thymidine, suggesting that impaired dTMP synthesis is indeed only a contributing mechanism of action for FCdR. Given the differential effects elicited by FCdR versus BVDU across pyrimidine nucleoside triphosphates, we also reasoned that putative interference with nucleic acid synthesis was unlikely a relevant mechanism to explain BVDU anticancer activity.

### DHFR is not the molecular target of brivudine

Despite inducing similar changes to several cellular components of purine and thymidylate metabolism, the conditional phenotypes for BVDU and MTX dramatically differed across cell lines. Therefore, we reasoned that DHFR was not a shared molecular target of MTX and BVDU-MP. To begin to investigate this, we sought to generate high-purity BVDU-MP. By uncoupling a UHPLC-MS system, we isolated BVDU-MP from reactions containing BVDU, ATP, and recombinant TK2, and then used a panel of NMP standards to estimate the concentration of pooled product (Figures 4H and S4I-K). We then developed a DHFR activity assay using LC-MS to measure NADP from reactions containing DHF (100 μM), NADPH (100 μM), and recombinant DHFR (Figures 4I and S4L). As anticipated, 10 μM MTX almost completely abolished NADP production, while equivalent addition of BVDU-MP to these reactions had negligible impact (Figure 4J).

Next, we asked whether BVDU and MTX differentially affect cellular folate profiles. When we treated K562 cells with doses that induced comparable growth defects in HPLM^+dS^, we found that each drug dramatically decreased the levels of folic acid (30-40%), 5-mTHF (10- to 20-fold), and 10-formyl-THF (60-80%); however, whereas THF levels were also reduced (40%) by MTX as would be expected, they were instead elevated by 60% in BVDU-treated cells (Figure 4K). These collective BVDU-induced changes resembled folate profiles previously described for cancer cells harboring genetic disruptions in the mitochondrial 1C pathway or a genetic knockout of *MTHFD1*, which encodes an enzyme in the cytosolic 1C pathway^77^. Thus, we speculated that the molecular target of BVDU treatment could be among this subset of proteins. Of note, when we profiled the same folates from K562 cells in HPLM^+dS^ containing RPMI-defined folic acid, we found that BVDU instead induced minimal effects versus vehicle (Figure S4M).

### Genome-wide CRISPR screens uncover genetic modifiers of brivudine sensitivity

To gain unbiased insights into the mechanism of BVDU cytotoxicity, we used a genome-wide single guide RNA (sgRNA) library to perform CRISPR screens in BVDU- and vehicle-treated K562 cells (Figure 5A). Following lentiviral infection and antibiotic selection in RPMI^+S^, we split and passaged cells in HPLM^+dS^ containing either BVDU at a concentration that elicited EC_25_ in a relevant culture flask format or DMSO (Figure S5A). For each gene, we calculated a scaled gene score and a probability of dependency in each screen after 15 population doublings as previously described (Table S4)^30^. Both screen datasets could discriminate core essential from nonessential genes^79^, and also contained a comparable number of essential genes (defined as probability of dependency > 0.5) (Figures S5B-C). We then standardized differential gene scores between the two screens. By integrating a Z-score cutoff of 3 with our probability of dependency data, we defined three categories of hits: resistance-conferring (positive; non-essential in both conditions), antagonizing (positive), and sensitizing (negative) (Figure 5B).

**Figure 5.**
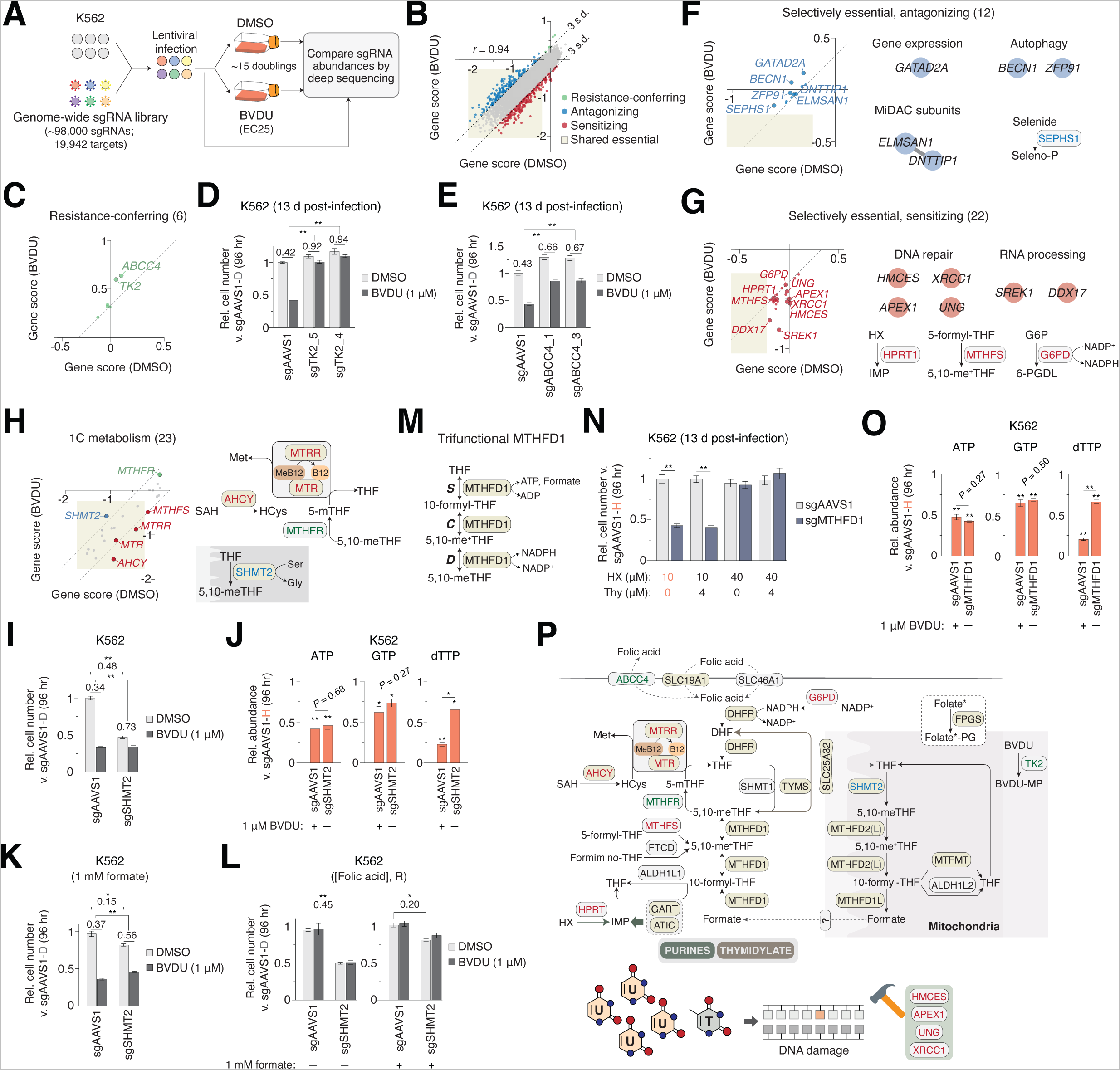
Genome-wide CRISPR screens for genetic modifiers of brivudine sensitivity See also Figure S5; Tables S3 and S4. (A) Schematic for genome-wide CRISPR modifier screens in K562 cells. (B) Comparison between DMSO- and BVDU-treated gene scores in the K562 modifier screens. Three defined sets of hits are resistance-conferring (green), antagonizing (blue), and sensitizing (red). Genes that scored as essential in both screens are in highlighted yellow box. *r*, Pearson’s correlation coefficient. (C) Resistance-conferring hits. (D-E) Relative growth of control and *TK2*-knockout (D) or *ABCC4*-knockout (E) cells treated with BVDU versus control cells in HPLM^+dS^ (mean ± SD, *n* = 3, \*\**P* < 0.005). (F-G) Selectively essential antagonizing (F) and sensitizing (G) hits. (H) Comparison between DMSO- and BVDU-treated gene scores across a panel of twenty-three genes involved in 1C metabolism (See Table S5). (I) Relative growth of control and *SHMT2*-knockout cells treated with BVDU versus control cells in HPLM^+dS^ (mean ± SD, *n* = 3, \*\**P* < 0.005). (J) Relative abundances of ATP, GTP and dTTP in BVDU-treated control and *SHMT2*-knockout cells versus control cells in HPLM^+dS^ (mean ± SD, *n* = 3, \*\**P* < 0.005, \**P* < 0.01). (K-L) Relative growth of control and *SHMT2*-knockout cells treated with BVDU versus control cells in HPLM^+dS^ (mean ± SD, *n* = 3, \*\**P* < 0.005, \**P* < 0.01). (M) Schematic of reactions catalyzed by tri-functional MTHFD1. S, 10-formyl-THF synthetase; C, 5,10-methynyl-THF cyclohydrolase; D, 5,10-meTHF dehydrogenase. (N) Relative growth of *MTHFD1*-knockout and control cells versus control cells in HPLM^+dS^ (mean ± SD, *n* = 3, \*\**P* < 0.005). (O) Relative abundances of ATP, GTP and dTTP in BVDU-treated control and *MTHFD1*-knockout cells versus control cells in HPLM^+dS^ (mean ± SD, *n* = 3, \*\**P* < 0.005). (P) Schematic of 1C metabolism with genes plotted in (H) and linked hit genes from (C) and (G). Screen hits and shared essential genes are colored as in (B) (top). BVDU-sensitizing hits involved in DNA repair (bottom). U, uracil; T, thymine. *Folylpolyglutamate synthase (FPGS) can convert several cellular folates to polyglutamylated derivatives.

Consistent with the notion that BVDU must be metabolically activated to elicit cytotoxicity, *TK2* was one of the top-two scoring resistance-conferring hits, along with *ABCC4* – a member of the ATP-binding cassette transporter family (Figure 5C). To complement our results showing that enforced *TK2* expression increases BVDU sensitivity in SEM cells, we engineered *TK2*-knockout K562 cells, which showed no baseline growth defects versus control cells but a nearly complete resistance to BVDU (Figure 5D). By engineering *ABCC4*-knockout K562 cells, we found that loss of *ABCC4* conferred a more modest 20-25% rescue of BVDU toxicity (Figure 5E). We considered the possibility that ABCC4 might mediate cellular import of BVDU but found that *ABCC4* deletion had little impact on the relative abundance of cellular BVDU (Figure S5D). However, prior uptake assays in insect cells demonstrated that ABCC4 could function as an efflux pump for folic acid^80^. Therefore, *ABCC4* knockout may alleviate BVDU sensitivity by increasing cellular folate levels.

Next, we identified antagonizing hits that were selectively essential in the vehicle screen, including genes involved in autophagy *(BECN1* and *ZFP91)*^81^, transcriptional repression, and selenophosphate synthesis (Figure 5F). Within this group, we also identified scaffolding subunits of the mitotic deacetylase complex, which are annotated as pairwise co-dependencies in DepMap as well^82–85^. Underlying causes that explain how loss of these genes protects cells against BVDU are not immediately apparent. In addition, we also identified sensitizing hits that were selectively essential in the BVDU screen, including several genes with roles in RNA processing or DNA repair *(HMCES, XRCC1, APEX1,* and *UNG*) (Figure 5G). Together, we reasoned that these hits were consistent with perturbations that could further sensitize cells with impaired de novo thymidylate synthesis. TYMS inhibition creates an imbalance of dUTP relative to dTTP that leads to increased misincorporation of uracil instead of thymine into DNA, which initiates base-excision repair (BER) mechanisms but can ultimately result in a “thymineless cell death”^40, 41, 86^. Uracil-DNA glycosylase (UNG) removes misincorporated uracil bases from DNA, leaving abasic sites that can be repaired by enzymes such as APEX1^40, 86^. Further, *XRCC1* encodes a molecular scaffold involved in BER and HMCES was recently characterized as a sensor that promotes the repair of abasic sites^87–89^. In addition, there is an increasing appreciation for the role of RNA processing in DNA repair^90, 91^. Notably, TYMS was not a BVDU modifier hit but scored as essential in both screens. This analysis further revealed *HPRT1* as a BVDU-essential hit, reflective of a greater dependence on the purine salvage pathway when de novo purine synthesis is impaired. Other BVDU-essential genes were involved in the pentose phosphate pathway *(G6PD)* and folate metabolism *(MTHFS)*. Previous work reported that the reaction catalyzed by glucose-6-phosphate dehydrogenase (G6PD) functions as a key source of the DHFR substrate NADPH in supporting folate metabolism, while others have suggested that 5,10-methynyl-THF synthetase (MTHFS) enhances purine production by effectively delivering 10-formyl-THF to enzymes in the de novo purine synthesis pathway^92–94^.

We also broadly examined CRISPR phenotypes for more than twenty genes with roles in 1C metabolism. Although most scored as essential in each screen, a small fraction was identified as BVDU-gene interaction hits as well (Figure 5H). Among these, sensitizing hits were linked to the methionine synthase (MTR) reaction that converts homocysteine and 5-mTHF to methionine and THF *(MTRR* and *AHCY)*, with *MTR* falling just below the set cutoff. We reason that loss of this reaction exacerbates BVDU cytotoxicity by “trapping” cellular folates as 5-mTHF, effectively depleting the availability of THF carriers that could otherwise be converted to de novo nucleotide synthesis substrates^95, 96^. We also identified *MTHFR* just below the scoring cutoff for resistance-conferring hits. Given that MTHFR irreversibly converts 5,10-meTHF to 5-mTHF, *MTHFR* deletion could alleviate BVDU sensitivity by effectively increasing the 5,10-meTHF pool available to TYMS. Interestingly, the only positive hit within this set of 1C pathway genes was *SHMT2*, which encodes the enzyme that initiates mitochondrial catabolism of serine to formate.

Since BVDU-induced effects on cellular folate levels resembled those reported in cancer cells with *SHMT2* knockdown among other genetic perturbations in the mitochondrial 1C pathway, we chose to further investigate this hit. We first created *SHMT2*-knockout K562 clonal cells, which showed a 50% growth defect versus control cells in HPLM^+dS^ but a 40% lower relative sensitivity to BVDU, confirming the BVDU-gene interaction and *SHMT2* essentiality suggested in our screen results (Figures 5I and S5E). However, whereas loss of *TK2* or *ABCC4* effectively reversed growth defects induced by BVDU, these seemingly differential treatment responses were instead linked to the baseline growth defect of *SHMT2*-knockout cells, establishing a key phenotypic distinction between resistance-conferring hits and antagonizing but shared essential genes. We then asked how loss of *SHMT2* affected the levels of purine and thymidylate metabolites following culture in HPLM^+dS^. *SHMT2* deletion decreased the levels of both ATP and GTP by an extent comparable to that in BVDU-treated control cells but elicited a more modest reduction in dTTP (Figure 5J). In addition, changes across de novo purine synthesis intermediates and dTMP were similar to those observed in BVDU-treated control cells, but *SHMT2* deletion caused an inverse effect on dUMP levels (Figure S5F). These results suggest that the BVDU modifier phenotype for *SHMT2* may be traced to partially redundant growth-impairing effects on nucleotide metabolism induced by either loss of *SHMT2* or BVDU treatment. Consistent with prior work in cells with disrupted mitochondrial 1C pathway genes^78, 97, 98^, increasing the formate levels in HPLM^+dS^ from 50 μM to 1 mM largely rescued the growth defects in *SHMT2*-knockout cells, but otherwise offered no protective effects to BVDU-treated control cells (Figure 5K). By contrast, adjusting folic acid availability to reflect RPMI-defined levels could not complement for the loss of *SHMT2* but reversed BVDU sensitivity regardless of *SHMT2* deletion or formate availability (Figure 5L). Together, these results revealed that 1C units are not limiting with BVDU treatment, suggesting that the target of BVDU-MP was unlikely a protein involved in the mitochondrial 1C pathway.

Since BVDU-induced changes to folate levels were similar to those reported for cells with deletion of the cytosolic *MTHFD1* as well, we next explored whether *MTHFD1*-knockout could recapitulate other BVDU treatment phenotypes – particularly given that *MTHFD1* was not a BVDU modifier hit but was essential in both screens. MTHFD1 is a tri-functional enzyme that catalyzes each of three sequential reactions, with activities conferred across two distinct domains: (1) N-terminal 5,10-methynyl-THF cyclohydrolase (C) and 5,10-meTHF dehydrogenase (D); and (2) C-terminal 10-formyl-THF synthetase (S) (Figure 5M)^99^. Using THF as a co-substrate, the forward MTHFD1-(S) reaction functions as a critical source of cytosolic 10-formyl-THF for de novo purine synthesis, as evidence has demonstrated that loss of *MTHFD1* can lead to purine auxotrophy^54,100^. Consistent with this, engineered *MTHFD1*-knockout K562 cells showed a 60% growth defect that was completely rescued by increasing HPLM-defined hypoxanthine levels by 4-fold but unaffected by the addition of 4 μM thymidine (Figures 5N and S5G). Similar to the case for *SHMT2* deletion, loss of *MTHFD1* decreased cellular ATP and GTP levels by extents reflective of those in BVDU-treated control cells but also caused a smaller relative reduction in dTTP (Figure 5O). Moreover, relative changes to de novo purine synthesis intermediates and TYMS reaction components more closely resembled those in *SHMT2*-knockout rather than BVDU-treated control cells, suggesting that loss of *MTHFD1* deletion could similarly phenocopy BVDU-induced effects on purine but not thymidylate metabolism (Figure S5H).

Collectively, these screen results supported the notion that TK2 mediates BVDU activation and that cellular folate levels affect BVDU sensitivity. We also identified several BVDU modifiers involved in either metabolic or DNA repair pathways that could be effectively linked to impairments of folate-dependent nucleotide synthesis (Figure 5P). Our follow-up data suggested the possibility that BVDU anticancer activity might be traced to multiple targets in cytosolic 1C metabolism.

### A dual-mechanism of action for brivudine and identification of other folic acid interactions

Given both the set of BVDU-sensitizing CRISPR hits with roles in BER pathways and that thymidine supplementation was necessary for the full rescue of BVDU sensitivity in HPLM^+dS^, we hypothesized that BVDU could affect human TYMS inhibition in cells lacking viral TK expression. To test this, we performed a single-dose cellular thermal shift assay (CETSA) in K562 cells after short-term treatment with BVDU or the established TYMS inhibitors MTX and FCdR. Indeed, each of these drugs could stabilize TYMS, and moreover, TYMS banding from FCdR-treated samples reflected the presence of both native protein and the expected ternary complex with 5,10-meTHF and FdUMP (Figures 6A and S6A). BVDU-treated samples showed similar banding but with the ratio instead favoring native versus bound TYMS, indicating that a fraction of cellular TYMS can perhaps form a similar ternary complex involving BVDU-MP in place of FdUMP. Collectively, our results suggest TYMS is one molecular target underlying the BVDU-folic acid interaction.

**Figure 6.**
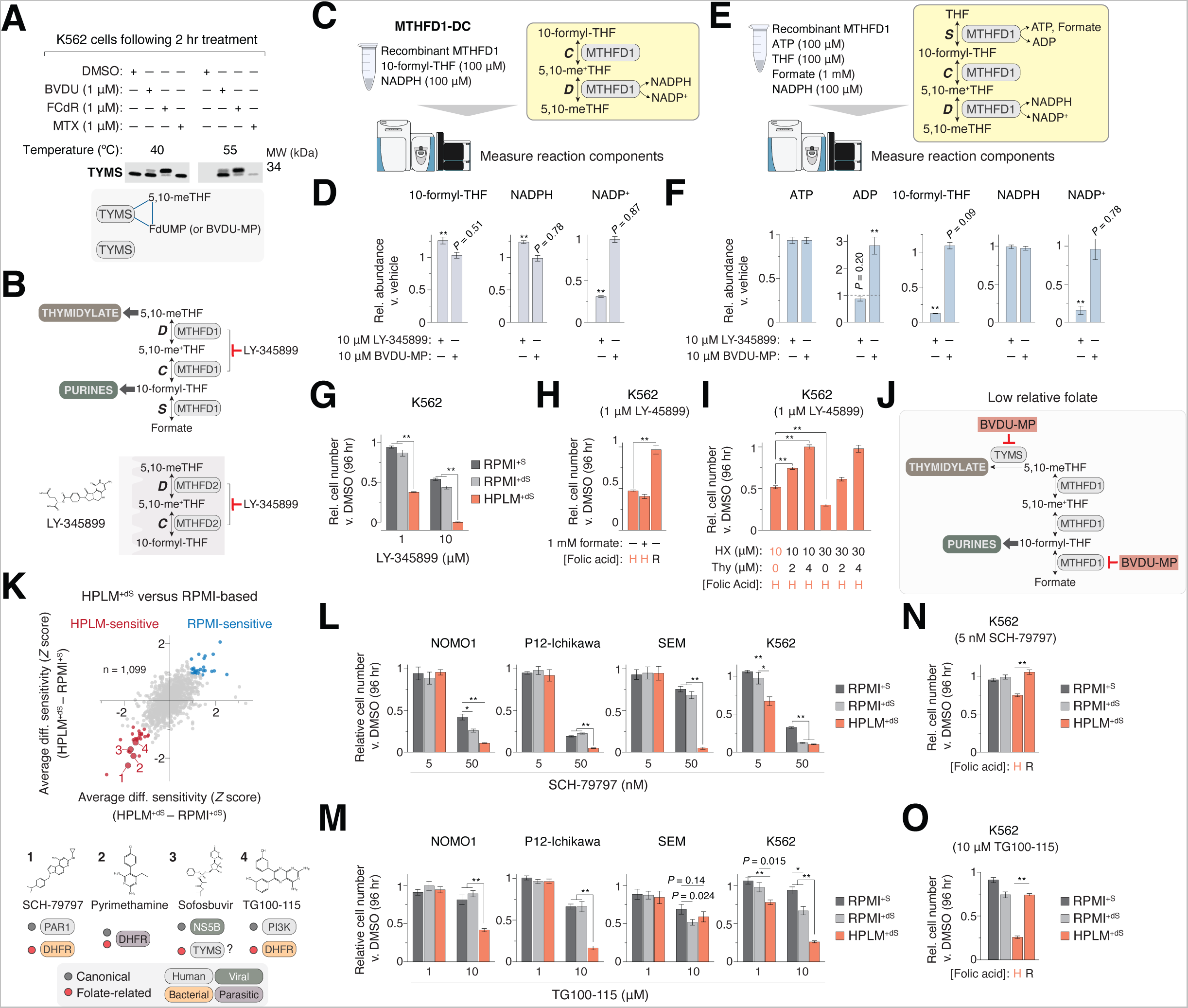
A dual-mechanism of action for brivudine and the identification of other drug-nutrient interactions with folic acid See also Figure S6; Table S1. (A) Immunoblot for expression of TYMS in cells treated with vehicle, BVDU, FCdR, or MTX in HPLM^+dS^ at indicated CETSA temperatures. M.W. standard is annotated (top). Covalent ternary complex formed between TYMS, 5,10-meTHF, and either FdUMP or BVDU-MP (bottom). (B) LY-345899 can inhibit the DC activities of cytosolic MTHFD1 and mitochondrial MTHFD2. (C) Schematic of an assay for MTHFD1-DC activity using LC-MS. (D) Relative abundances of MTHFD1-DC reaction components versus vehicle to the assay in (C) (mean ± SD, *n* = 3, \*\**P* < 0.005). (E) Schematic of an assay for trifunctional MTHFD1 activity using LC-MS. (F) Relative abundances of MTHFD1 reaction components to the assay in (E) (mean ± SD, *n* = 3, \*\**P* < 0.005). (G-I) Relative growth of cells treated with LY-345899 versus DMSO (mean ± SD, *n* = 3, \*\**P* < 0.005). H, HPLM-defined concentration. R, RPMI-defined concentration. (J) Proposed model for conditional BVDU cytotoxicity. In low folate conditions, BVDU-MP disrupts de novo purine and thymidylate synthesis through inhibition of MTHFD1 and TYMS, respectively. (K) Comparison between averaged HPLM^+dS^-RPMI^+dS^ and HPLM^+dS^-RPMI^+S^ phenotypes across the three screened lines (top). Other HPLM-sensitive hits with putative links to folate metabolism are highlighted (bottom). (L-O) Relative growth of cells treated with SCH-79797 (L and N) or TG100-115 (M and O) versus DMSO (mean ± SD, *n* = 3, \*\**P* < 0.005, \**P* < 0.01).

Next, we considered that MTHFD1 could be a distinct target of BVDU-MP, providing a link to the BVDU-mediated inhibition of de novo purine synthesis revealed by our phenotypic profiling and screen results. To investigate this, we first sought to develop LC-MS-based MTHFD1 activity assays, reasoning that these could be used to delineate domain-specific activities in reactions containing recombinant MTHFD1 (Figure S6B). Previous studies have described compounds that inhibit the D/C activities of MTHFD1 or the mitochondrial MTHFD2, including LY-345899 – a folate analog that displays a 7-fold greater affinity for MTHFD1 (Figure 6B)^101–103^. Consistent with this, addition of 10 μM LY-345899 to reactions containing the two substrates specific to MTHFD1-DC – 10-formyl-THF (100 μM) and NADPH (100 μM) – reduced NADP production by 70%, whereas equivalent BVDU-MP had little effect (Figures 6C-D). When we then evaluated reactions instead containing substrates specific to forward MTHFD1-S activity – ATP (100 μM), THF (100 μM), and formate (1 mM) – we detected negligible product formation (10-formyl-THF and ADP). However, supplementing these reactions with NADPH (100 μM), the final substrate input across MTHFD1 domains, enabled the detection of each MTHFD1-S product and NADP (Figures 6E and S6C-D). Using this tri-functional MTHFD1 activity assay, we found that LY-345899 not only impaired NADP production but also markedly reduced 10-formyl-THF levels, whereas BVDU-MP increased ADP levels by nearly 3-fold but did not otherwise affect the other detected reaction components (Figure 6F). Moreover, equivalent addition of either BVDU or MTX in this assay did not impact the levels of any MTHFD1 reaction components (Figure S6E). Interestingly, prior work with monofunctional MTHFD1-S orthologs from prokaryotes has suggested that the synthetase reaction proceeds in sequential steps, whereby formation of a formylphosphate intermediate is followed by the release of ADP and formylation of THF^104,105^. This mechanism effectively uncouples the synthesis of ADP and 10-formyl-THF. Collectively, we reason that LY-345899 inhibits MTHFD1-DC as anticipated, while BVDU-MP interferes with the MTHFD1-S domain that generates cytosolic 10-formyl-THF for de novo purine synthesis. Of note, overall effects observed upon the addition of either drug in this tri-functional MTHFD1 activity assay are difficult to completely reconcile given the reversibility of each reaction.

Notably, when we tested LY-345899 against K562 cells, we observed 40-50% stronger growth defects in HPLM^+dS^ versus both RPMI-based media, which could be rescued by increasing exogenous folic acid to RPMI-defined levels but not by supplementing with 1 mM formate (Figures 6G-H). These results reveal that increasing folic acid availability can also relieve inhibition of the MTHFD1-DC domain and further suggest that MTHFD2 is unlikely a relevant target of LY-345899 in K562 cells. Given that MTHFD1-DC is downstream of MTHFD1-S in the cytosolic branch of 1C metabolism, we hypothesized that the LY345899-folic acid interaction was specific to a disruption of de novo dTMP synthesis. Indeed, thymidine supplementation completely rescued LY-345899 sensitivity, while boosting hypoxanthine levels by 4-fold surprisingly exacerbated growth defects, consistent with prior work that examined how these nutrients affect cancer cell responses to either LY-345899 or an alternative MTHFD1-DC inhibitor (Figure 6I)^106,107^.

Collectively, we propose a dual-mechanism model to explain conditional BVDU anticancer activity. BVDU is converted to BVDU-MP by mitochondrial TK2 regardless of folic acid availability. Upon export to the cytosol, BVDU-MP effectively competes with folates to mediate a hierarchical inhibition of MTHFD1 and TYMS, ultimately disrupting de novo purine and thymidylate synthesis, respectively (Figure 6J). While prior work using in vitro assays showed that BVDU-MP could inhibit viral or human homologs of TYMS^108^, this effector metabolite shares little similarity with reported MTHFD1/2 inhibitors^101,109^, which are also otherwise typically characterized for activities against MTHFD1-DC. Moreover, this mechanism of BVDU action is unique compared to those described for other pyrimidine nucleoside analogs, including FCdR and additional dS-sensitive hits revealed by our chemical screens.

Next, we identified other HPLM-sensitive compounds that we speculated could similarly form interactions with folic acid based on their reported links to DHFR orthologs or similarity to pyrimidine nucleoside analogs (Figure 6K). Among these were SCH-79797, a PAR-1 antagonist that was recently characterized as a bacterial DHFR inhibitor, and TG100-115, a PI3K inhibitor recently identified as a bacterial DHFR-binding compound as well^110,111^. Given their shared non-canonical target, we chose to investigate these two hits. When we tested these drugs against our cell line panel, we largely confirmed their HPLM-sensitive phenotypes, though TG100-115 did not elicit medium-dependent responses in SEM cells (Figures 6L-M and S6F-G). Consistent with our rationale, HPLM^+dS^ with RPMI-defined levels of folic acid could normalize K562 cell responses to each compound relative to RPMI^+dS^ (Figures 6N-O). Interestingly, TG100-115 was among the 2% weakest scoring compounds in our RPMI^+dS^-RPMI^+S^ profile, while SCH-79797 was instead within the top 15% strongest scoring (Figure S6H). Of note, when we evaluated a more recent lot of the commercial solution initially used to incorporate RPMI vitamins into HPLM, we found that working concentrations for biotin and folic acid were more comparable to those in basal RPMI, but still 2-fold lower than their respective RPMI-defined values (Figure S6I).

### Identification of other conditional anticancer compounds

We could also use our data to identify compounds with conditional phenotypes likely linked to the availability other nutrients. For example, the top-three scoring RPMI-sensitive hits were (1) CB-839, a glutaminase (GLS) inhibitor that has been tested in cancer patients^112^; (2) deguelin, a natural product shown to inhibit cancer cell growth through putative inhibition of PI3K^113^; and (3) apilimod, a reported PIKFYVE inhibitor being tested in lymphoma patients (Figure 7A)^114–116^. Our previous genome-wide and secondary CRISPR screens in K562 cells revealed *GLS* among the RPMI-essential hits, a conditional dependency that we traced to differential pyruvate availability (Figure 7B)^30^. Surprisingly, this CRISPR phenotype for *GLS* was specific to RPMI^+dS^ despite our detection of only sub-physiologic pyruvate in RPMI^+S^. Nonetheless, we previously phenocopied conditional *GLS* essentiality by treating K562 cells with CB-839 in RPMI^+dS^ versus HPLM^+dS^. When we tested CB-839 against our cell line panel, we found that responses in K562 and NOMO1 cells were each reflective of the conditional CRISPR phenotype for *GLS* in K562 cells, while those in the other two lines were instead consistent with suggested RPMI-sensitive phenotypes regardless of FBS dialysis (Figures 7C and S7A). These results not only support the notion that cell sensitivity to GLS inhibition is dictated by combined cell-intrinsic and environmental factors, but also suggest that co-targeting systemic pyruvate levels may improve CB-839 efficacy in some cases. Of note, while prior work in other cell lines traced conditional CB-839 toxicity to cystine availability^117^, the levels of cysteine/cystine in HPLM and RPMI differ by less than two-fold (Figure S7B).

**Figure 7.**
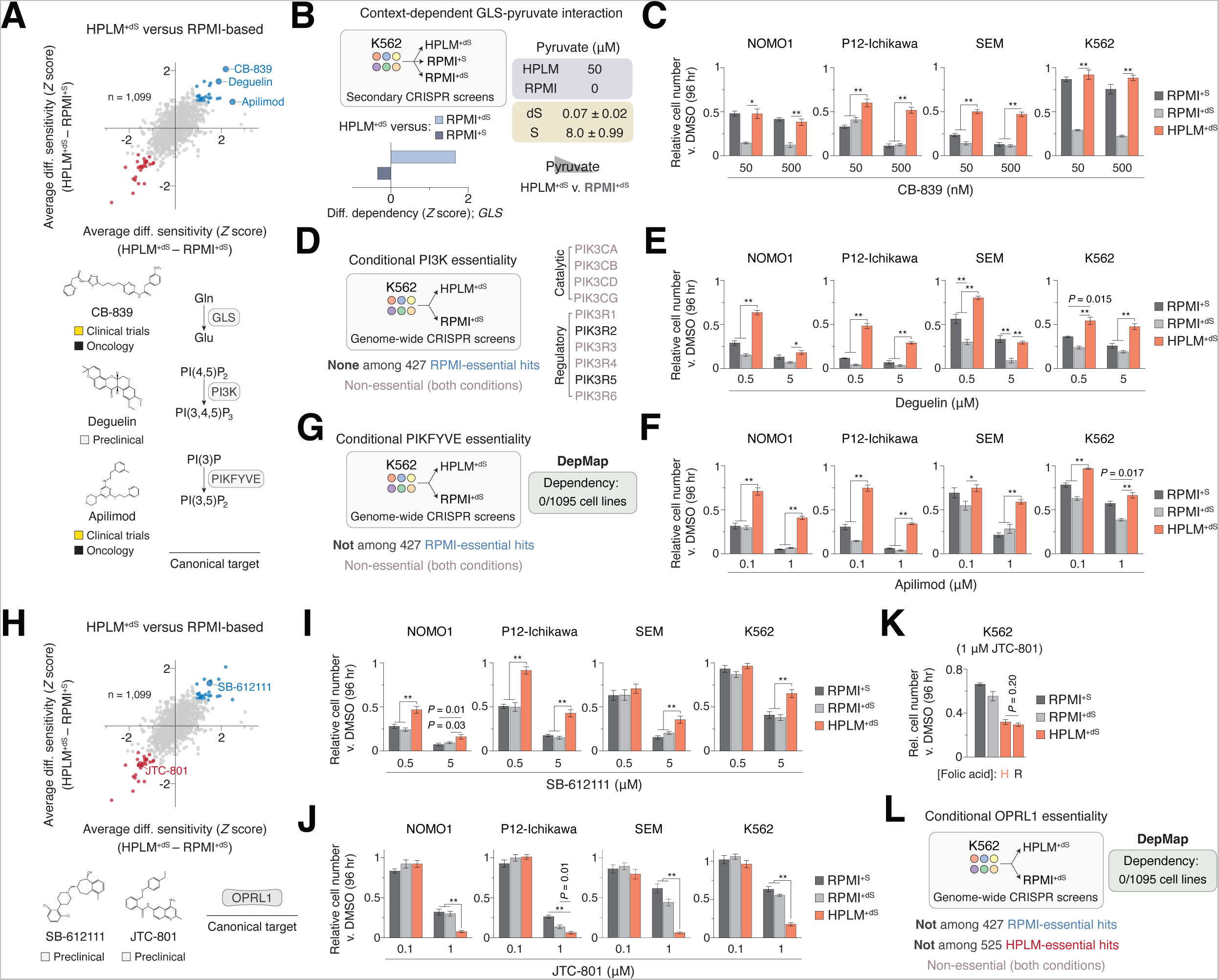
Identification of other conditional anticancer compounds See also Figure S7; Table S1. (A) Comparison between averaged HPLM^+dS^-RPMI^+dS^ and HPLM^+dS^-RPMI^+S^ phenotypes. Top-three scoring RPMI-sensitive hits are labeled (top). Canonical targets of each compound (bottom). (B) Conditional CRISPR phenotypes for *GLS* from secondary screens in K562 cells (left). Defined pyruvate levels in HPLM and RPMI. Concentrations of pyruvate in 10% FBS (dS, dialyzed; S, untreated) (mean ± SD, *n* = 3) (right). (C) Relative growth of cells treated with CB-839 versus DMSO (mean ± SD, *n* = 3, \*\**P* < 0.005, \**P* < 0.01). (D) Data for catalytic *(PIK3C)* and regulatory *(PIK3R)* PI3K subunits from genome-wide CRISPR screens in K562 cells. (E-F) Relative growth of cells treated with deguelin (E) or apilimod (F) versus DMSO (mean ± SD, *n* = 3, \*\**P* < 0.005, \**P* < 0.01). (G) *PIKFYVE* data from genome-wide CRISPR screens in K562 cells and DepMap. (H) Comparison between averaged HPLM^+dS^-RPMI^+dS^ and HPLM^+dS^-RPMI^+S^ phenotypes. Two OPRL1 antagonists are labeled. (I-K) Relative growth of cells treated with SB-612111 (I) and JTC-801 (J-K) versus DMSO (mean ± SD, *n* = 3, \*\**P* < 0.005). (L) *OPRL1* data from genome-wide CRISPR screens in K562 cells and DepMap.

Although deguelin is annotated as a PI3K inhibitor in the MIPE library, we did not identify any catalytic *(PIK3C)* or regulatory *(PIK3R)* PI3K subunits among the roughly 400 RPMI-essential genes in our previous genome-wide CRISPR screens (Figure 7D)^30^. Nonetheless, we observed RPMI-sensitive phenotypes for deguelin across our cell line panel, though relative growth defects in RPMI^+S^ were more modest in SEM cells (Figures 7E and S7C). It is interesting to consider that other putative targets and pathways have been reported to explain deguelin cytotoxicity as well^113^. We also validated the conditional phenotype for apilimod across cell lines (Figures 7F and S7D). Despite existing evidence for PIKFYVE-apilimod binding^114, 118^, *PIKFYVE* was nonessential in our prior genome-wide CRISPR screens and also absent from the set of RPMI-essential hits (Figure 7G). Remarkably, *PIKFYVE* is also not annotated as a genetic dependency in any of the ∼1,100 cancer cell lines from DepMap^85^. Together, these results demonstrate that deguelin and apilimod form drug-nutrient interactions that are not immediately apparent, and further suggest that both compounds can mediate anticancer effects through non-canonical mechanisms of action^119^.

Our analysis also revealed two hit compounds each annotated as opioid receptor (OPRL1) antagonists that otherwise showed inverse conditional phenotypes: JTC-801 (HPLM-sensitive); and SB-612111 (RPMI-sensitive) (Figure 7H)^120, 121^. Indeed, we confirmed both phenotypes for at least one dose in each cell line, and in addition, found that HPLM^+dS^ with RPMI-defined folic acid could not rescue JTC-801-induced growth defects in K562 cells (Figures 7I, 7J, 7K, and S7E-F). Notably, *OPRL1* was a nonessential gene that was also absent from the combined set of nearly 1,000 conditionally essential hits in our genome-wide CRISPR screens and is also not annotated as a dependency in any cell lines from DepMap (Figure 7L). Moreover, RNA-seq data indicate that *OPRL1* shows negligible expression in more than 90% of the 1,400 human cancer cell lines evaluated, including three from our panel (Figure S7G)^122^. These results further demonstrate that conditional phenotypes for compounds with otherwise common putative targets can vary, and that non-oncology drugs can elicit anticancer effects^21^.

## CONCLUSIONS

Here we have demonstrated that chemical screens in HPLM versus traditional media can reveal drug treatment responses that depend on nutrient availability. Compounds with conditional anticancer activities include investigational compounds and drugs tested in patients or approved for use in the treatment of cancer and a variety of non-oncology indications. Conditional treatment phenotypes can also vary with natural cell-intrinsic diversity even among a panel of only four cell lines. This suggests that high-throughput screens in HPLM should facilitate the identification of new drug candidates for a variety of cancer cell types under nutrient conditions that more faithfully model those faced in the human body. The ability to screen large numbers of compounds against various cell types in HPLM, including libraries comprised only of approved agents, should further make it possible to uncover drug repurposing opportunities for cancer and other diseases. Based on recently described methods to identify protein-metabolite interactions^123, 124^, it is also interesting to consider that target-based chemical screens in HPLM could reveal binding ligands that would perhaps be masked in conventional media.

Conditional phenotypes for several chemotherapeutics were traced to nucleotide salvage pathway substrates defined only in HPLM versus RPMI or that are otherwise provided by the serum supplement used in most complete media. Using a systematic approach, we also identified an unforeseen drug-nutrient interaction between folic acid and BVDU, an antiviral agent approved to treat herpes zoster in several countries. By integrating drug-nutrient complementation assays, metabolomics, CRISPR-based BVDU modifier screens, metabolite synthesis, and in vitro enzyme assays, we characterized a unique dual-mechanism for conditional BVDU sensitivity. TK2 first converts BVDU to the active effector metabolite BVDU-MP, which in turn disrupts de novo purine and thymidylate synthesis in low folate conditions by inhibiting the cytosolic 1C pathway enzymes MTHFD1 and TYMS. Of note, we further demonstrate a strategies to identify selectively essential genetic modifiers of drug sensitivity and to distinguish resistance-conferring genetic deletions from other antagonizing drug-gene interactions.

Previously identified conditional phenotypes for 5-FU and CB-839 were each more broadly recapitulated across cell lines, but our results reveal that RPMI-sensitive phenotypes for CB-839 can differ between cell lines for RPMI^+S^ and RPMI^+dS^. This highlights the complexity in translating preclinical drug sensitivities on the basis of phenotypes that vary not only with cell-intrinsic factors but also with the basal and serum components of culture media. We also confirm additional drug-nutrient interactions each traced to folic acid for two compounds (SCH-79797 and TG100-115) with distinct canonical targets each unrelated to 1C metabolism but that otherwise share recently described links to bacterial DHFR. In addition, we validate conditional phenotypes for apilimod, a drug being tested in cancer patients, and a number of investigational compounds (deguelin, SB-612111, and JTC-801) as well. By leveraging our previous conditional gene essentiality profiles in K562 cells, we find that canonical targets reported for these compounds are encoded by genes that not only lack an equivalent (conditional) CRISPR phenotype, but in multiple cases, have also not been annotated as essential in any of the nearly 1,100 cell lines from DepMap. This suggests that systematic approaches will be needed to identify drug-nutrient interactions formed by these compounds as well.

Conditional phenotypes from chemical screens can be traced to drug-nutrient interactions in human cells. We show that this strategy offers the potential to explore mechanisms of action by integrating comparisons of chemical and genetic perturbations with others between drug- and gene-nutrient interactions. This also raises the interesting possibility to exploit conditional lethality by developing treatment strategies that combine molecular therapeutics with dietary or enzyme-mediated modulation of systemic metabolites^31–33^.

## ACKNOWLEDGEMENTS

We thank L. Chen for assistance with chemical screen data processing, K. Huggler for upkeep of the LC-MS system, members of the Cantor lab for discussions, E. Frenkel and the Whitehead Institute Functional Genomics Platform for kindly providing the human sgRNA library, and K. Tharp, E. Bresnick, and C. Alexander for manuscript comments. We also thank D.M. Sabatini for helping to facilitate the collaboration that initiated this study. This work was supported by the NIH (K22CA225864 to J.R.C., T32GM145470 to N.J.R., and T32GM135066 to A.L.H.) and startup funds from the Morgridge Institute for Research.

## AUTHOR CONTRIBUTIONS

M.D.H. and J.R.C. conceived the study. M.D.H., K.M.W., and J.R.C. designed the drug screens. J.R.C. designed the remainder of the study. K.M.W. and T.D.L. optimized and performed chemical screens. N.J.R. and K.M.W. analyzed chemical screen data. J.R.C. curated clinical development phase and indication data for library compounds with assistance from K.M.W. J.R.C. performed most of experiments with assistance from K.M.F., A.L.H., and N.J.R. K.M.F. optimized and performed folate profiling. A.L.H. optimized and performed most of immunoblots. J.R.C. analyzed and interpreted the experimental data with assistance from K.M.F. J.R.C. performed the CRISPR screens. N.J.R. analyzed the CRISPR screen data. J.R.C. wrote the manuscript. All authors discussed the manuscript. J.R.C. supervised the studies.

## SUPPLEMENTAL INFORMATION

Supplemental information includes all methods, seven figures, and five tables.

## SUPPLEMENTAL INFORMATION

### Supplemental figure legends

**Figure S1.**
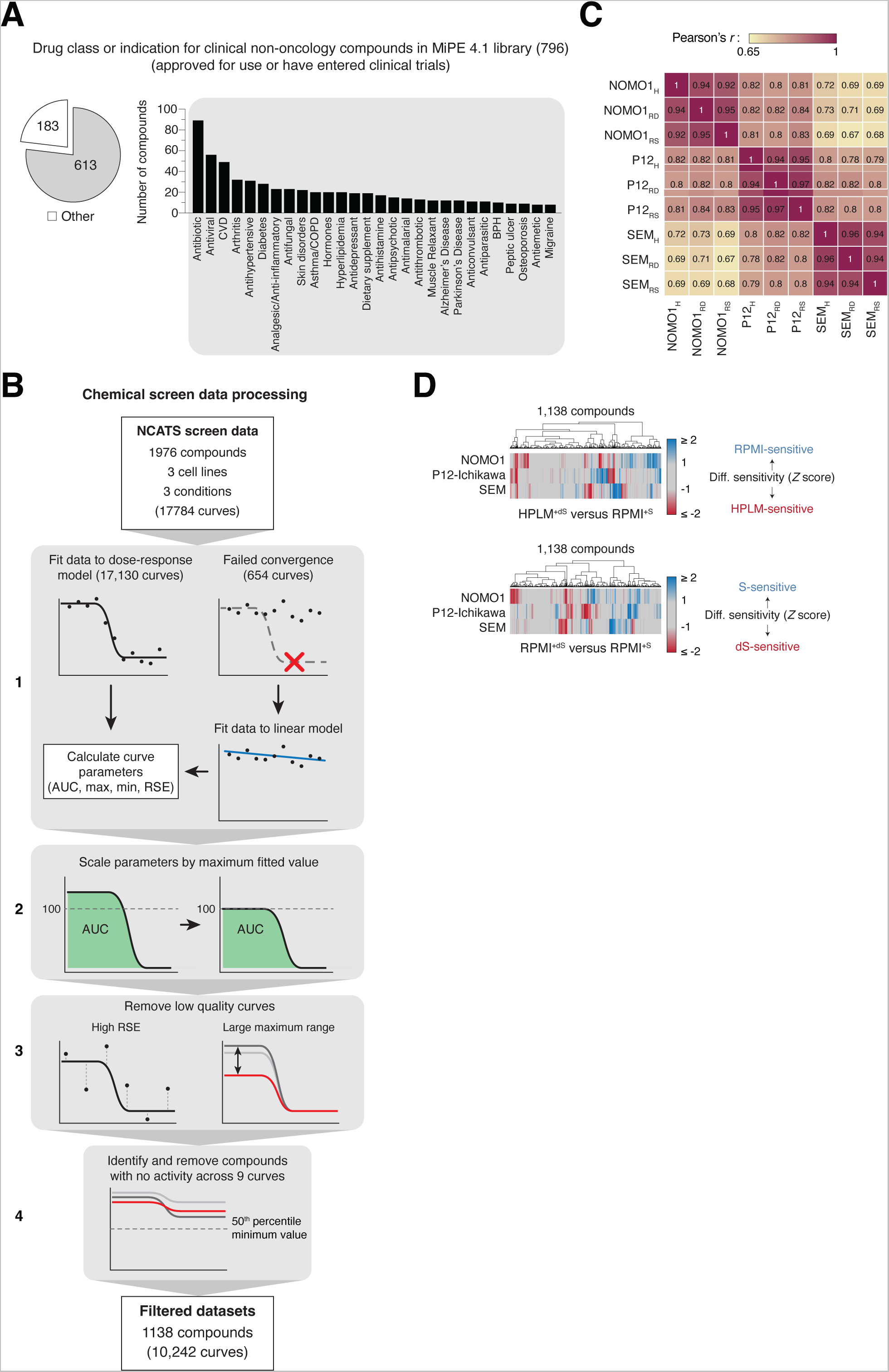
Chemical screen analysis, Related to Figure 1. (A) Composition of non-oncology compounds in MIPE 4.1 library by manually curated drug class or indication, among those either approved for use in humans or that have entered clinical trials. CVD, cardiovascular disease; COPD, chronic obstructive pulmonary disease; BPH, benign prostatic hyperplasia. (B) Data processing workflow. See **Quantification and Statistical Analysis** for additional detail. *(1)* Normalized viability data for 17,784 total dose-response curves were fit to a 4-parameter log-logistic model. 654 curves failed to converge and were fit using linear regression. *(2)* Curves with a maximum viability greater than 100% were scaled by the maximum fitted value. *(3)* Curves with residual standard error (RSE) values above cell line-dependent 98^th^ percentiles were removed (left); Cell line-specific curves with greater than a 15% difference in maximum viability between two or more conditions for a given compound were removed (right). *(4)* Compounds with minimum values greater than the respective median values in each of the screen datasets were removed. Together, 1,138 filtered compounds were shared across all cell line-medium combinations. AUC, area under the curve. (C) Response score correlations between nine chemical screens spanning three cell lines and three conditions. H, HPLM^+dS^; RD, RPMI^+dS^; RS, RPMI^+S^. (D) Cluster maps showing HPLM^+dS^ versus RPMI^+S^ (top) and RPMI^+dS^ versus RPMI^+S^ (bottom) conditional phenotypes in three cell lines.

**Figure S2.**
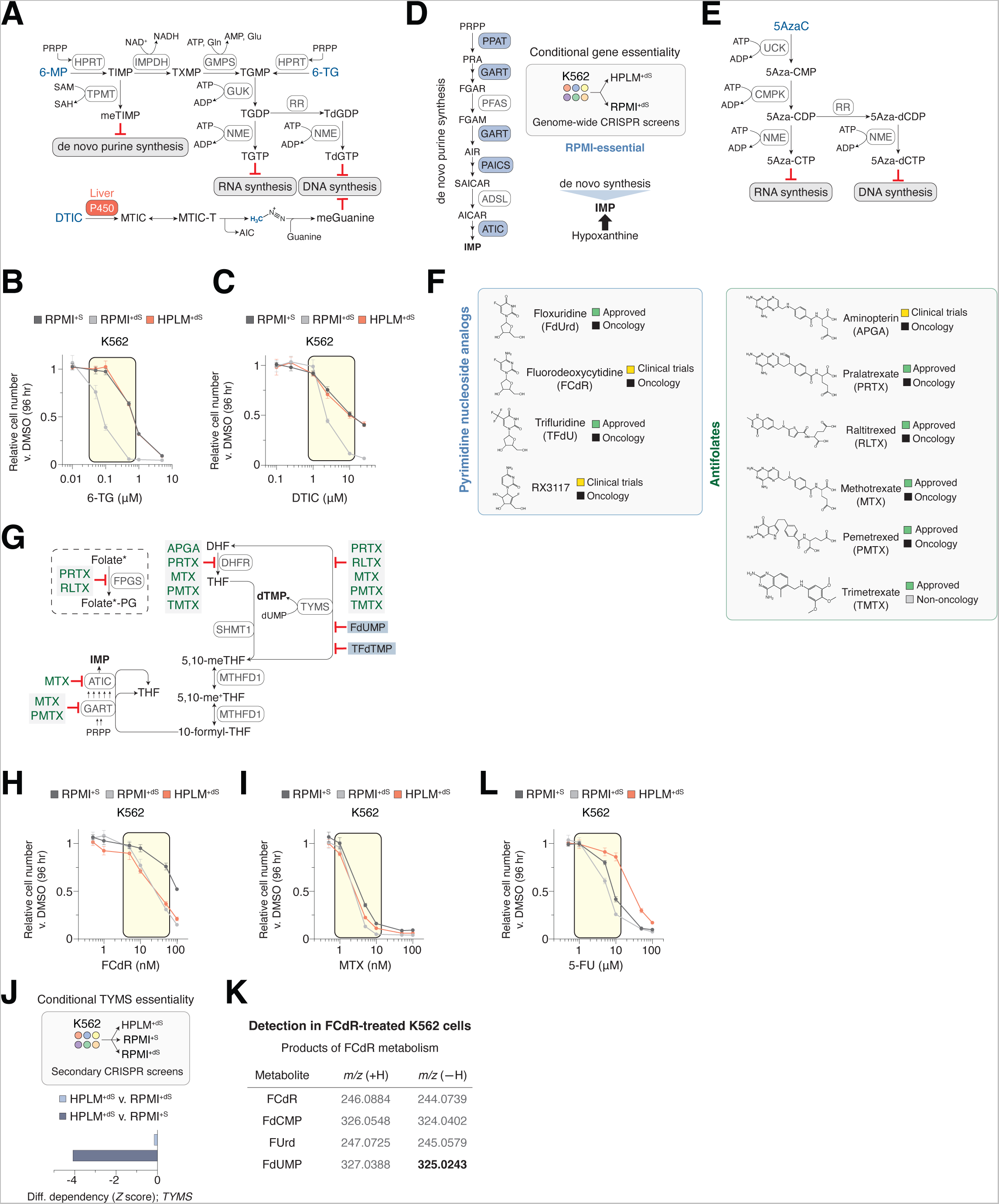
Additional data related to drug-nutrient interactions with nucleotide salvage pathway substrates, Related to Figure 2. (A) Cellular conversion of RPMI-sensitive purine analogs to metabolites that mediate cytotoxic effects. The canonical mechanism of dacarbazine (DTIC) activity involves an initial activation step catalyzed by liver-specific P450. 6-MP, 6-Mercaptopurine; 6-TG, 6-Thioguanine. (B-C) Dose-responses of K562 cells to 6-TG (B) and DTIC (C) (mean ± SD, *n* = 3). Concentration range spanned for two dose-responses tested across the remaining three cell lines (yellow box). (D) Schematic for the de novo purine synthesis pathway. Enzymes encoded by genes that were identified as RPMI-essential hits from previous genome-wide CIRSPR screens in K562 cells (shaded blue). Hypoxanthine is a salvage pathway substrate that can be used to generate IMP. (E) Cellular conversion of RPMI-sensitive 5AzaC to metabolites that mediate cytotoxic effects. (F) dS-sensitive pyrimidine nucleoside analogs (left) and antifolates (right). (G) Schematic of folate-dependent enzyme targets that may be directly or indirectly inhibited by dS-sensitive antifolates. *Folylpolyglutamate synthase (FPGS) can convert several cellular folates to polyglutamylated derivatives. Fluorodeoxyuridine monophosphate (FdUMP) and trifluoromethyl deoxyuridine monophosphate (TFdTMP) are effector derivatives of FdUrd/FCdR and TFdU that can inhibit TYMS as well. (H-I) Dose-responses of K562 cells to FCdR (H) and MTX (I) (mean ± SD, *n* = 3). Concentration range spanned for two dose-responses tested across the remaining three cell lines (yellow box). (J) Conditional CRISPR phenotypes for *TYMS* from secondary screens in K562 cells. (K) Mass-to-charge ratios (m/z) for various products of cellular FCdR metabolism based on either the addition (+H) or removal (-H) of a proton adduct (left). Peaks corresponding to FdUMP in negative ionization mode (-H) were the only ones from this table that could be detected in FCdR- treated K562 cells. (L) Dose-responses of K562 cells to 5-FU (mean ± SD, *n* = 3). Concentration range spanned for two dose-responses tested across the remaining three cell lines (yellow box).

**Figure S3.**
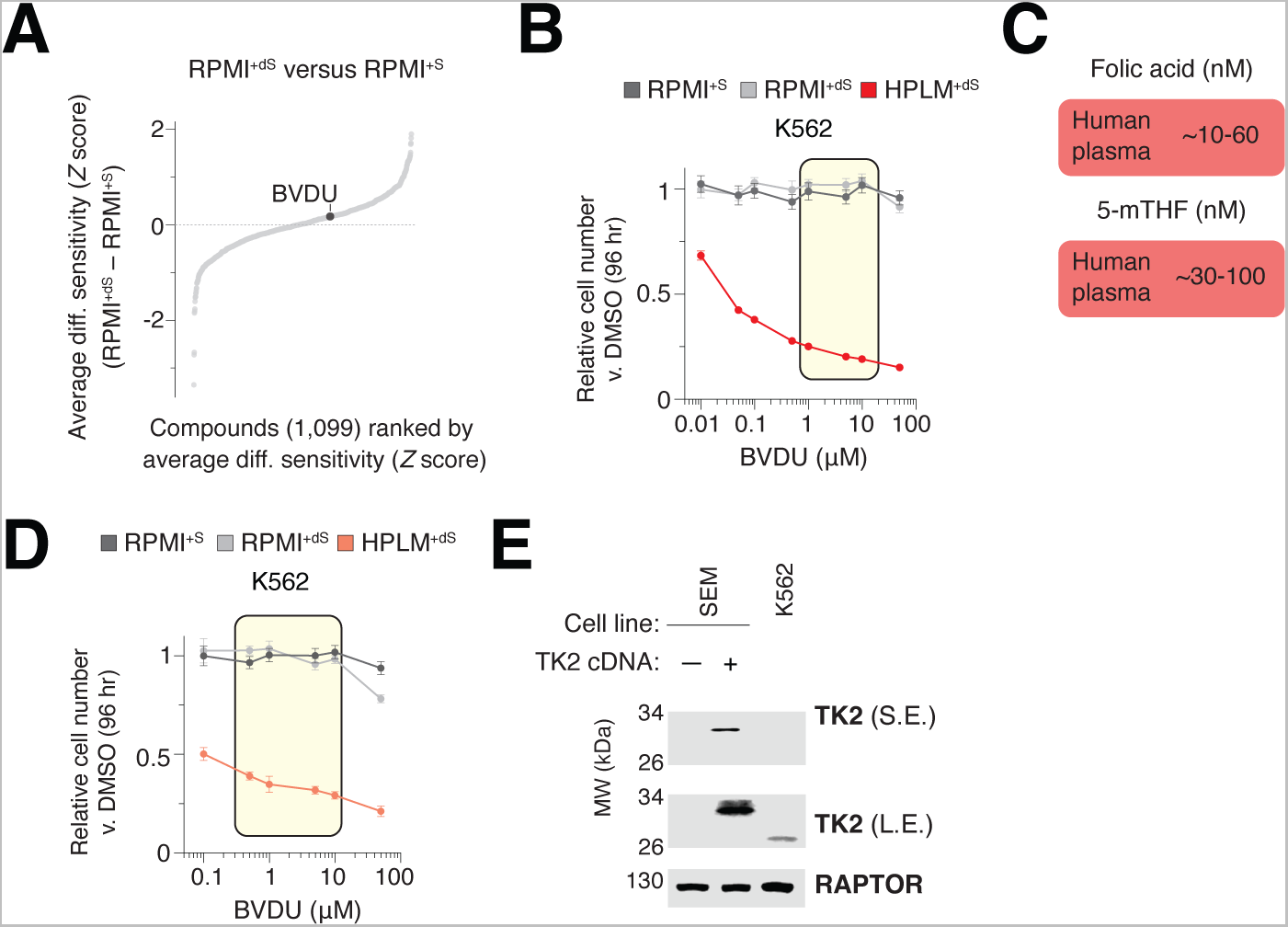
Additional data related to identification of a drug-nutrient interaction between brivudine and folic acid, Related to Figure 3. (A) Compounds ranked by average RPMI^+dS^-RPMI^+S^ sensitivity across three cell lines. Brivudine (BVDU) is labeled. (B) Dose-responses of K562 cells to BVDU (mean ± SD, *n* = 3). Concentration range spanned for two dose-responses tested across the remaining three cell lines (yellow box). (C) Reported concentration ranges for folic acid and 5-methyl-THF (5-mTHF) in human plasma^70–73^. (D) Dose-responses of K562 cells to BVDU (mean ± SD, *n* = 3). Concentration range spanned for two dose-responses tested across the remaining three cell lines (yellow box). (E) Immunoblot for expression of TK2. M.W. standards are annotated. RAPTOR served as the loading control. S.E., short exposure. L.E., long exposure.

**Figure S4.**
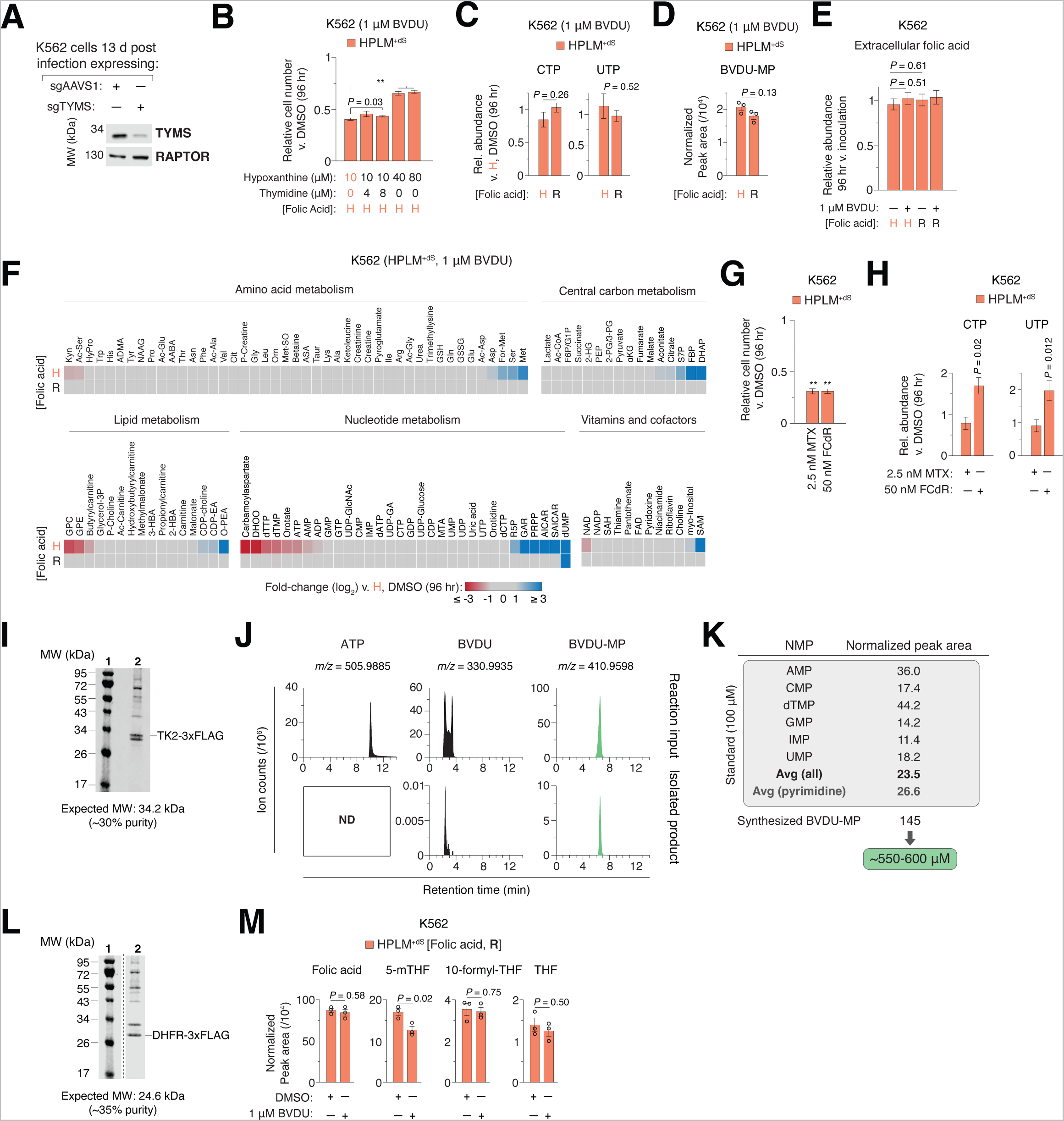
Additional data related to brivudine interferes with folate-dependent nucleotide synthesis, Related to Figure 4. (A) Immunoblot for expression of TYMS. M.W. standards are annotated. RAPTOR served as the loading control. TYMS band intensities differ by ∼5-fold between the two samples. (B) Relative growth of cells treated with BVDU versus DMSO (mean ± SD, *n* = 3, \*\**P* < 0.005). H, HPLM-defined concentration (450 nM). (C) Relative abundances of CTP and UTP in BVDU-treated and control cells versus control cells in HPLM^+dS^ (mean ± SD, *n* = 3, \*\**P* < 0.005). R, RPMI-defined concentration (2.27 μM). (D) Cellular abundances of BVDU-MP following BVDU treatment in HPLM^+dS^ (mean ± SD, *n* = 3). (E) Relative abundances of folic acid in HPLM^+dS^ following 96 hr treatment with BVDU versus those at inoculation (mean ± SD, *n* = 3). (F) Heatmap of relative metabolite abundances in cells treated with BVDU in HPLM^+dS^ containing HPLM- (top) or RPLMI-defined (bottom) folic acid versus control cells in HPLM^+dS^. Metabolite clusters are each sorted by log_2_-transformed fold change of the top row (n = 3). Metabolite abbreviations can be found in Table S3. (G) Relative growth of cells treated with FCdR or MTX versus DMSO (mean ± SD, *n* = 3, \*\**P* < 0.005). (H) Relative abundances of CTP and UTP in FCdR- and MTX-treated versus control cells in HPLM^+dS^ (mean ± SD, *n* = 3, \*\**P* < 0.005). (I) Pseudocolor Coomassie-stained gel imaged using a LICOR Odyssey FC. 1: M.W. standards, 2: TK2-3xFLAG. (J) Extracted ion chromatograms at mass-to-charge (m/z) ratios, each in negative ionization mode, for ATP, BVDU, and BVDU monophosphate (BVDU-MP) at indicated retention times for samples extracted from reactions containing purified recombinant TK2 with ATP and BVDU (top) and the isolated BVDU-MP (bottom). See **Method Details**. (K) Normalized peak areas across a panel of NMP chemical standards and the in vitro synthesized BVDU-MP. A concentration for the stock BVDU-MP was estimated based on the average of these standard areas – with little effect on this average if considering only the pyrimidine NMPs. (L) Pseudocolor Coomassie-stained gel imaged using a LICOR Odyssey FC. 1: M.W. standards, 2: DHFR-3xFLAG. (M) Abundances of folate metabolites in BVDU-treated and control cells in HPLM^+dS^ with RPMI- defined folic acid (mean ± SD, *n* = 3).

**Figure S5.**
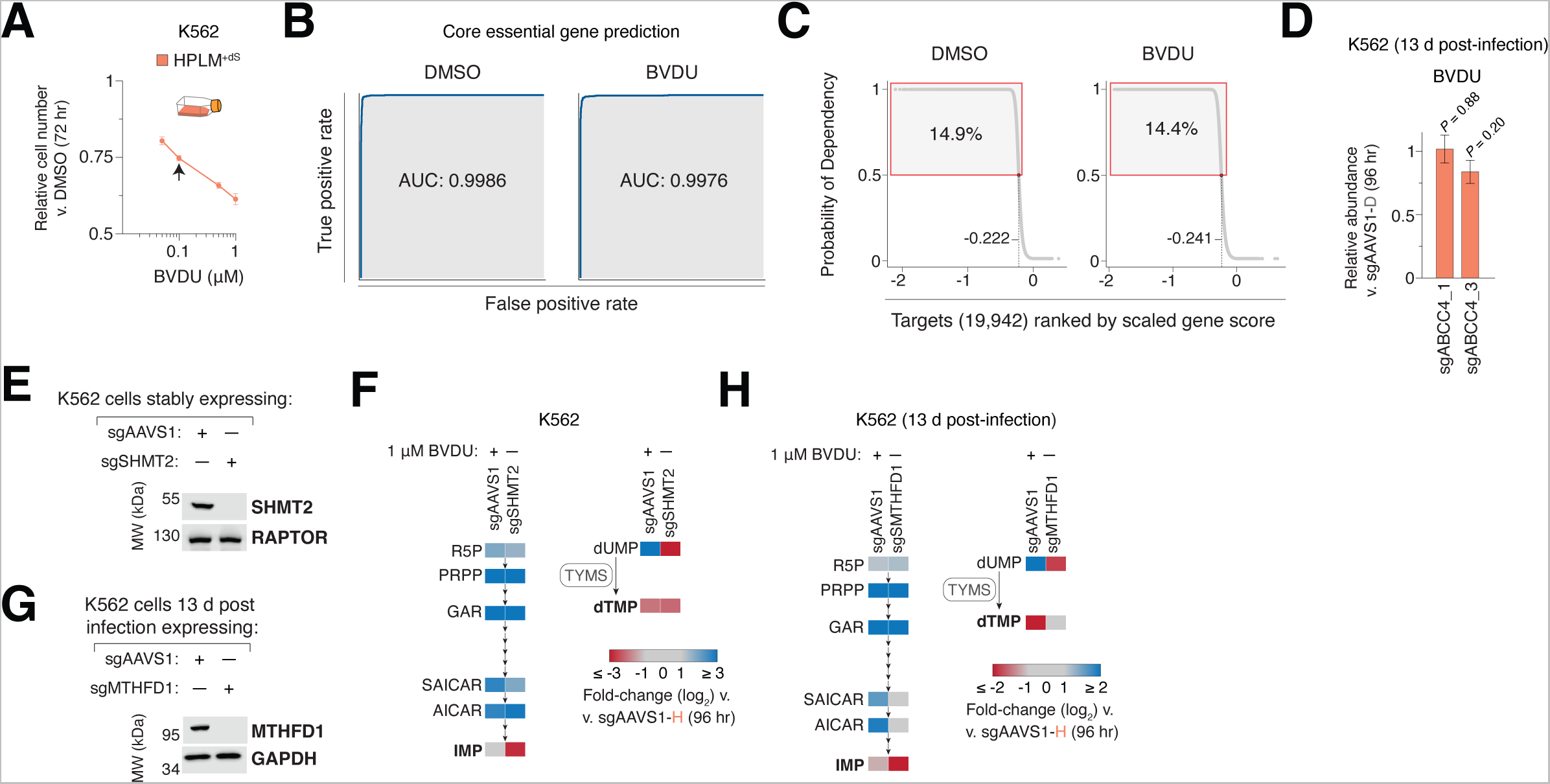
Additional data related to genome-wide CRISPR screens for genetic modifiers of brivudine sensitivity, Related to Figure 5. (A) Relative growth of cells treated with BVDU versus DMSO in T-25 flasks (mean ± SD, *n* = 3). Arrow indicates the dose that elicited a ∼25% growth defect. (B) Receiver operator characteristic (ROC) curves for the prediction of core essential genes using datasets from BVDU modifier CRISPR screens. (C) Plots of library targets ranked by probability of dependency from genome-wide K562 screens in HPLM^+dS^ with DMSO vehicle (left) or BVDU (right). Red box indicates probability > 0.5. Dashed lines mark gene scores at the threshold for gene essentiality in each screen. (D) Relative abundance of BVDU in *ABCC4*-knockout versus control cells following treatment with BVDU in in HPLM^+dS^ (mean ± SD, *n* = 3). (E) Immunoblot for expression of SHMT2. M.W. standards are annotated. RAPTOR served as the loading control. (F) Heatmap of relative abundances for metabolites in the de novo purine synthesis pathway (left) and for two TYMS reaction components (right). BVDU-treated control and *SHMT2*-knockout cells versus control cells HPLM^+dS^. (G) Immunoblot for expression of MTHFD1. M.W. standards are annotated. GAPDH served as the loading control. (H) Heatmap of relative abundances for metabolites in the de novo purine synthesis pathway (left) and for two TYMS reaction components (right). BVDU-treated control and *MTHFD1*-knockout cells versus control cells HPLM^+dS^.

**Figure S6.**
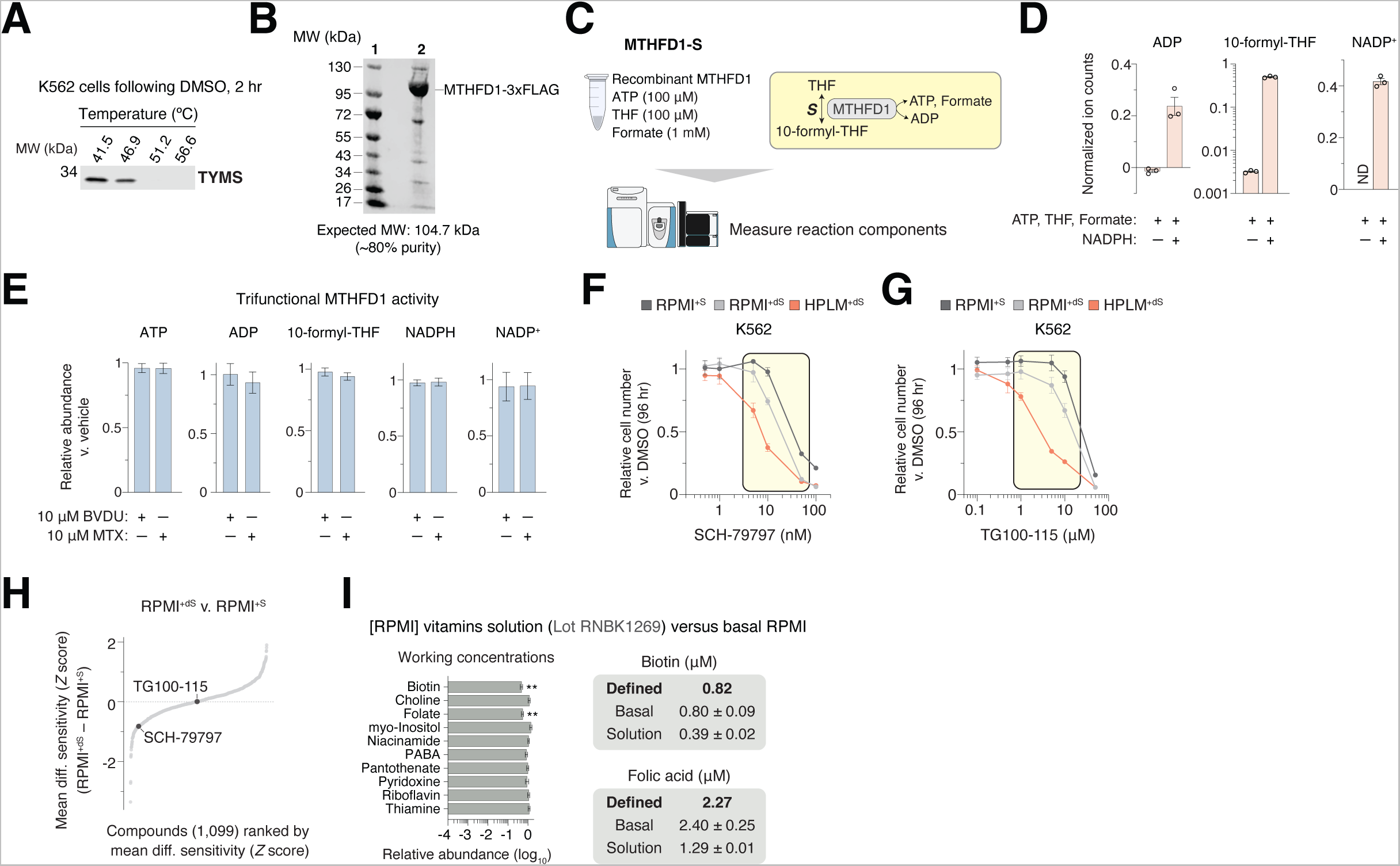
Additional data related to a dual-mechanism of action for brivudine and the identification of other drug-nutrient interactions with folic acid, Related to Figure 6. (A) Immunoblot for expression of TYMS in cells treated with vehicle in HPLM^+dS^ across indicated CETSA temperatures. M.W. standard is annotated. (B) Pseudocolor Coomassie-stained gel imaged using a LICOR Odyssey FC. 1: M.W. standards, 2: MTHFD1-3xFLAG. (C) Schematic of an assay for MTHFD1-S activity using LC-MS. (D) Normalized ion counts for ADP, 10-formyl-THF, and NADP in both the MTHFD1-S (-NADPH) and trifunctional MTHFD1 (+NADPH) assays (mean ± SD, *n* = 3). Correction for background ADP resulted in slightly negative values reflective of noise in the MTHFD1-S assay. (E) Relative abundances of MTHFD1 reaction components following addition of BVDU or MTX versus vehicle to the assay in (E) (mean ± SD, *n* = 3, \*\**P* < 0.005). (F-G) Dose-responses of K562 cells to SCH-79797 (F) and TG100-115 (G) (mean ± SD, *n* = 3). Concentration range spanned for two dose-responses tested across the remaining three cell lines (yellow box). (H) Compounds ranked by average RPMI^+dS^-RPMI^+S^ sensitivity across three cell lines. SCH-79797 and TG100-115 are labeled. (I) Relative working concentrations of vitamins in RPMI 100X vitamin solution (Sigma R7256, Lot RNBK1269) versus basal RPMI (Thermo 11879, Lot 2458379) (mean ± SD, *n* = 3, \*\**P* < 0.005) (left). Defined and working concentrations of biotin and folic acid (mean ± SD, *n* = 3) (right). RPMI also contains 3.69 nM vitamin B12, which could not be detected by the profiling method.

**Figure S7.**
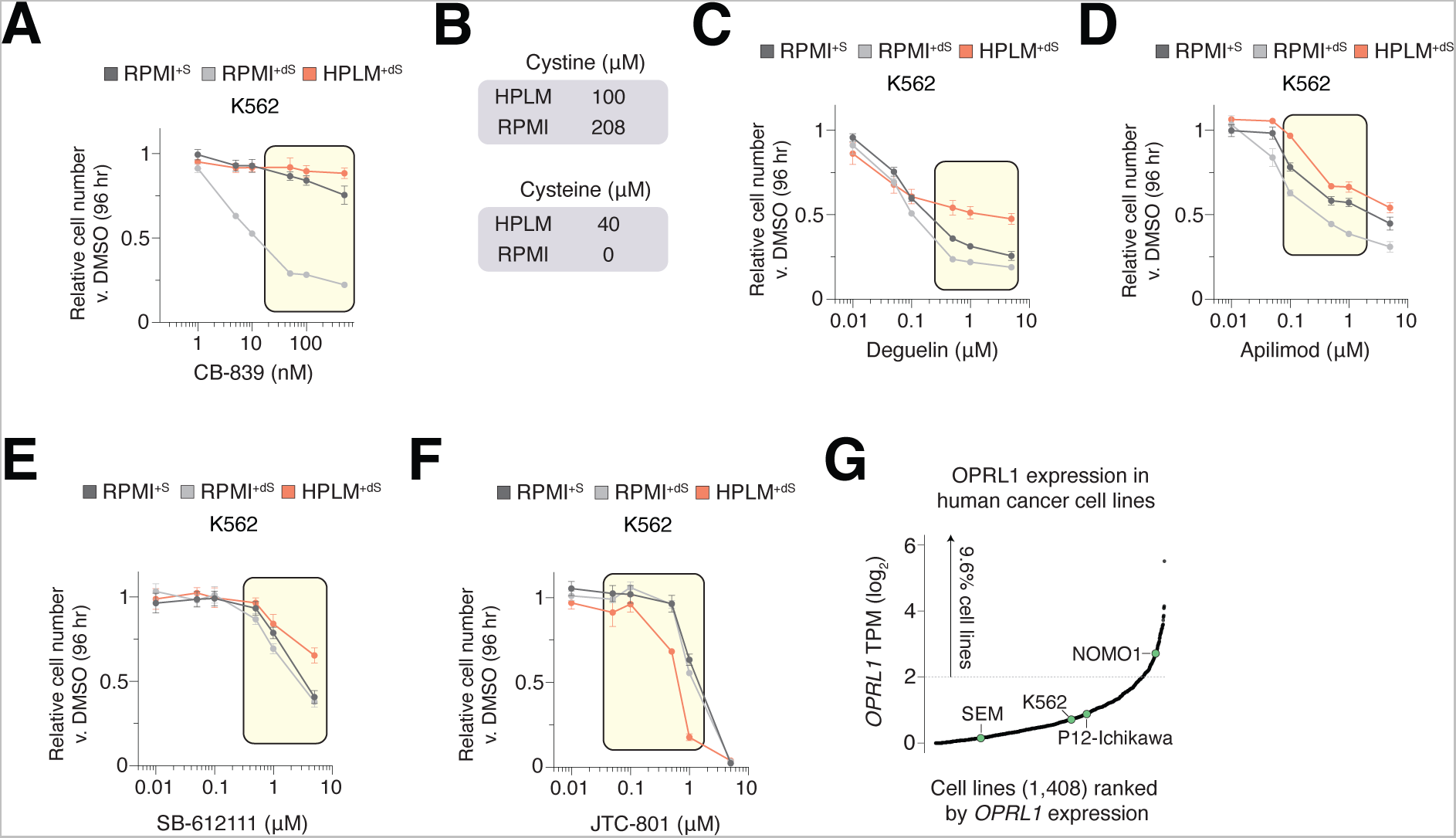
Additional data related to identification of other conditionally anticancer compounds, Related to Figure 7. (A) Dose-responses of K562 cells to CB-839 (mean ± SD, *n* = 3). Concentration range spanned for two dose-responses tested across the remaining three cell lines (yellow box). (B) Defined cystine and cysteine levels in HPLM and RPMI. (C-F) Dose-responses of K562 cells to deguelin (C), apilimod (D), SB-612111 (E), or JTC-801 (F) (mean ± SD, *n* = 3). Concentration range spanned for two dose-responses tested across the remaining three cell lines (yellow box). (G) Human cancer lines ranked by *OPRL1* RNA levels from RNA-seq data in DepMap. Labeled points indicate cell lines in this study. TPM, transcripts per million.

### Supplemental Tables

**Table S1: Datasets related to chemical screens, Related to** **Figure 1**

**Table S2: Synthetic media construction, Related to** **Figure 3**

**Table S3: Datasets related to metabolite profiling, Related to Figures 3-5**

**Table S4: Datasets related to CRISPR screens, Related to** **Figure 5**

**Table S5: Oligonucleotides used in this study, Related to Methods**

## REAGENTS AND RESOURCES

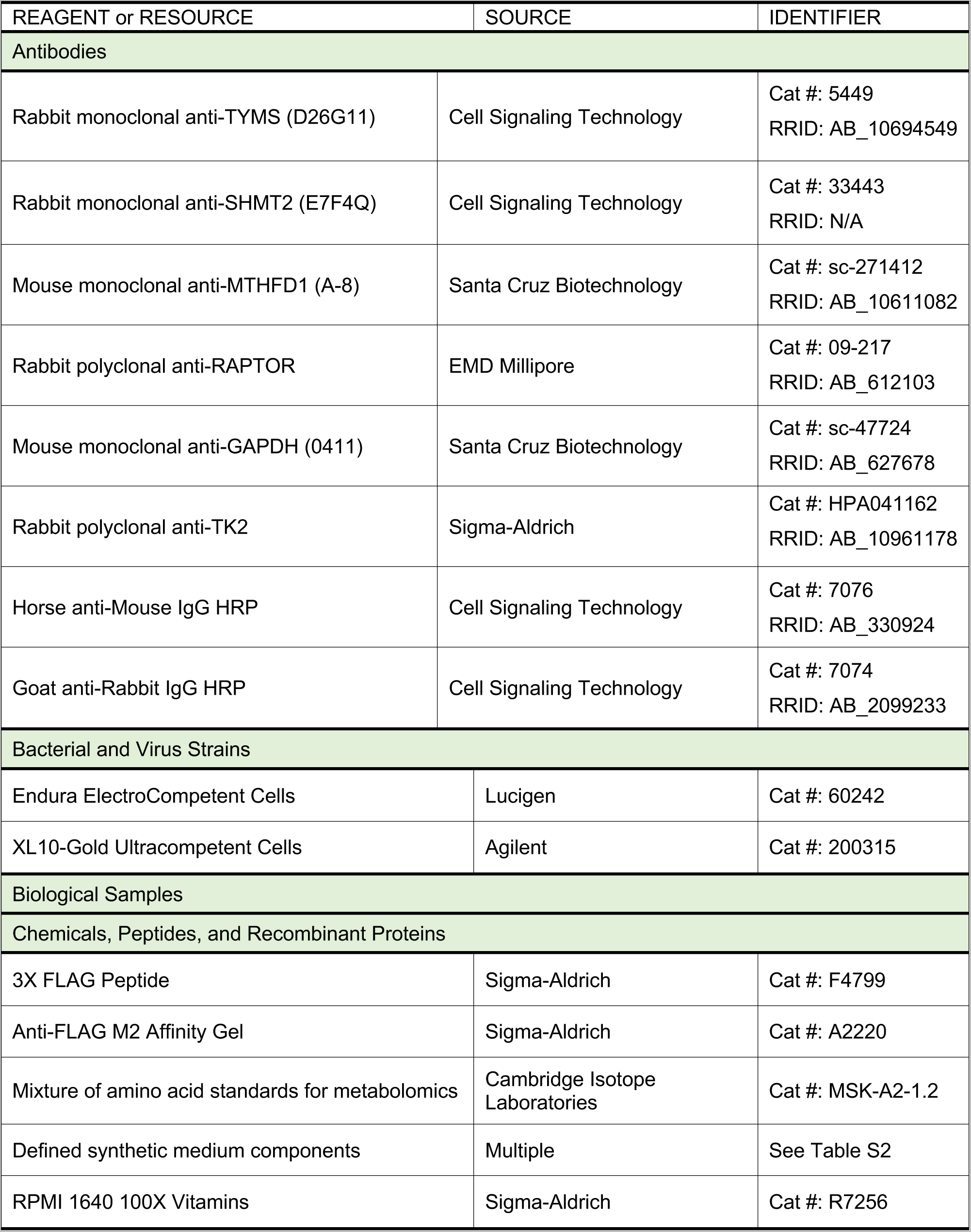
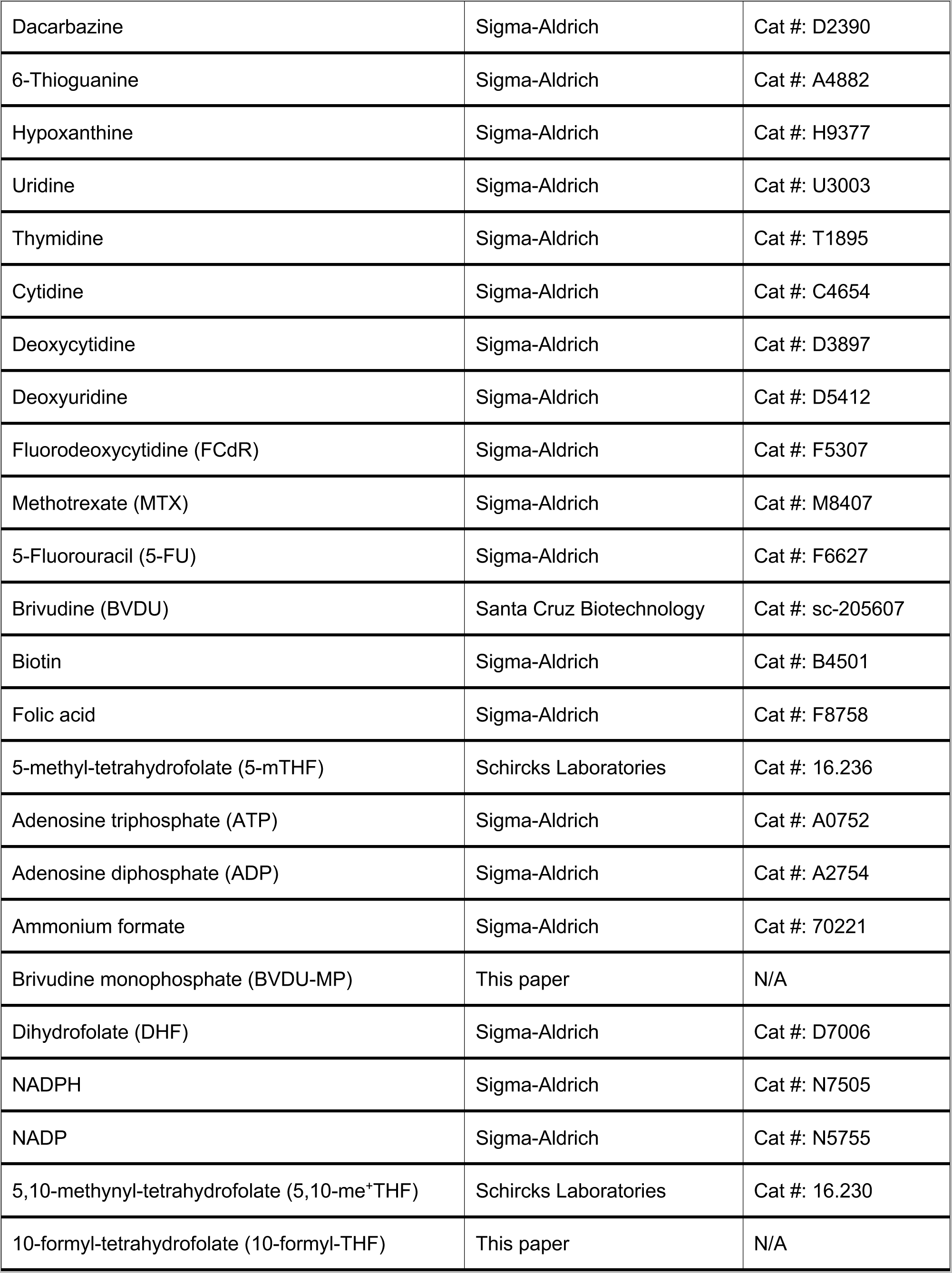
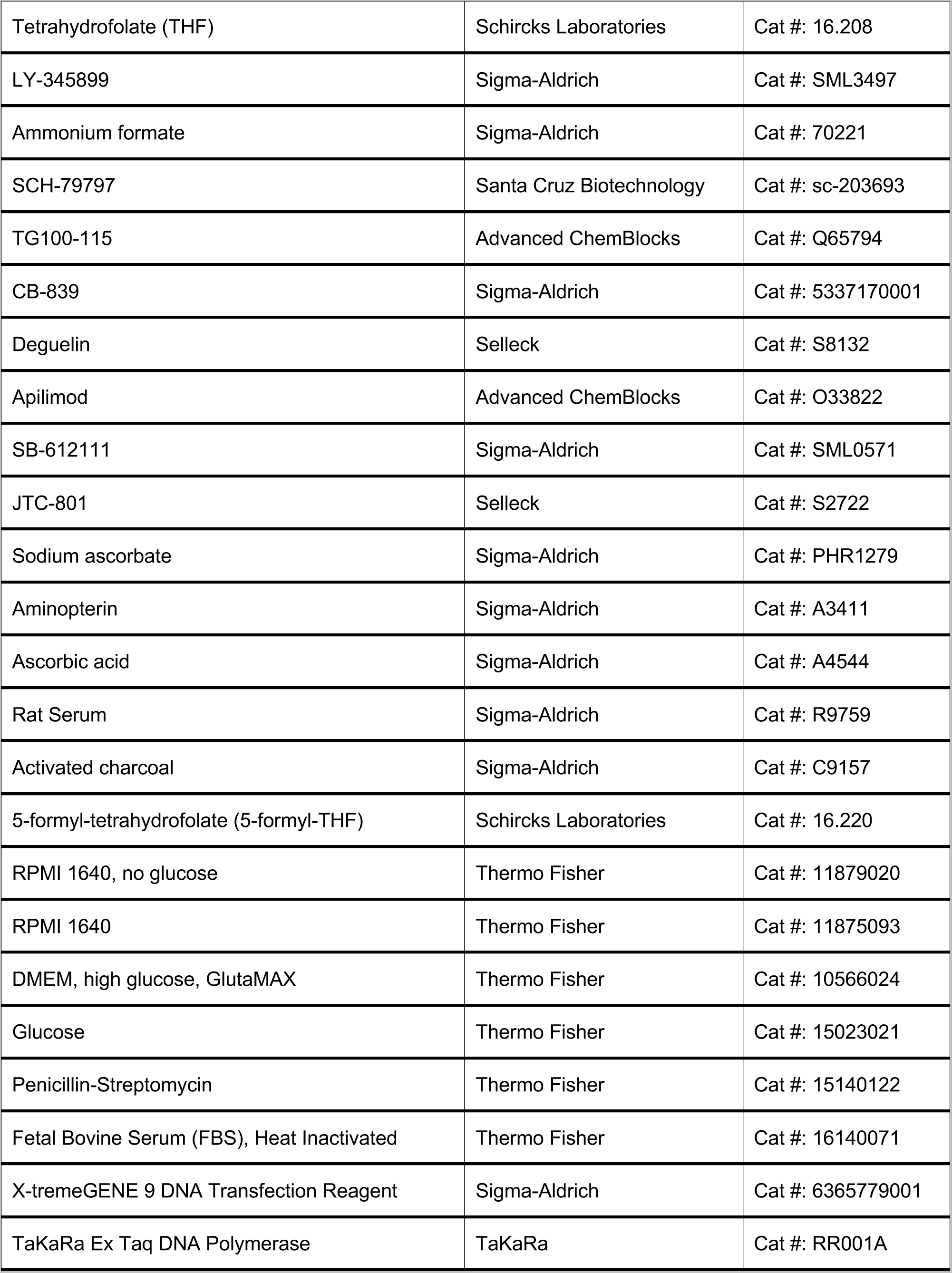
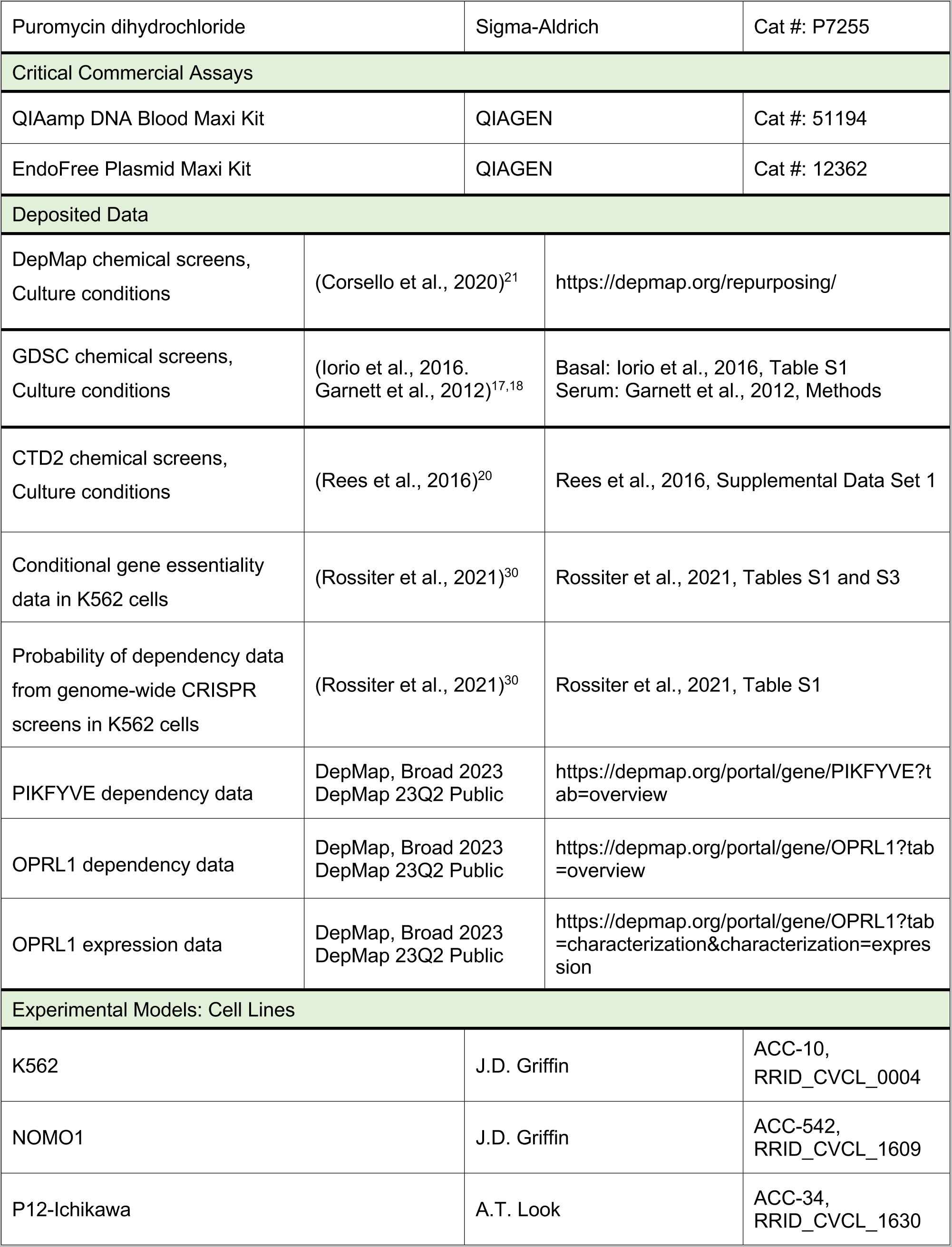
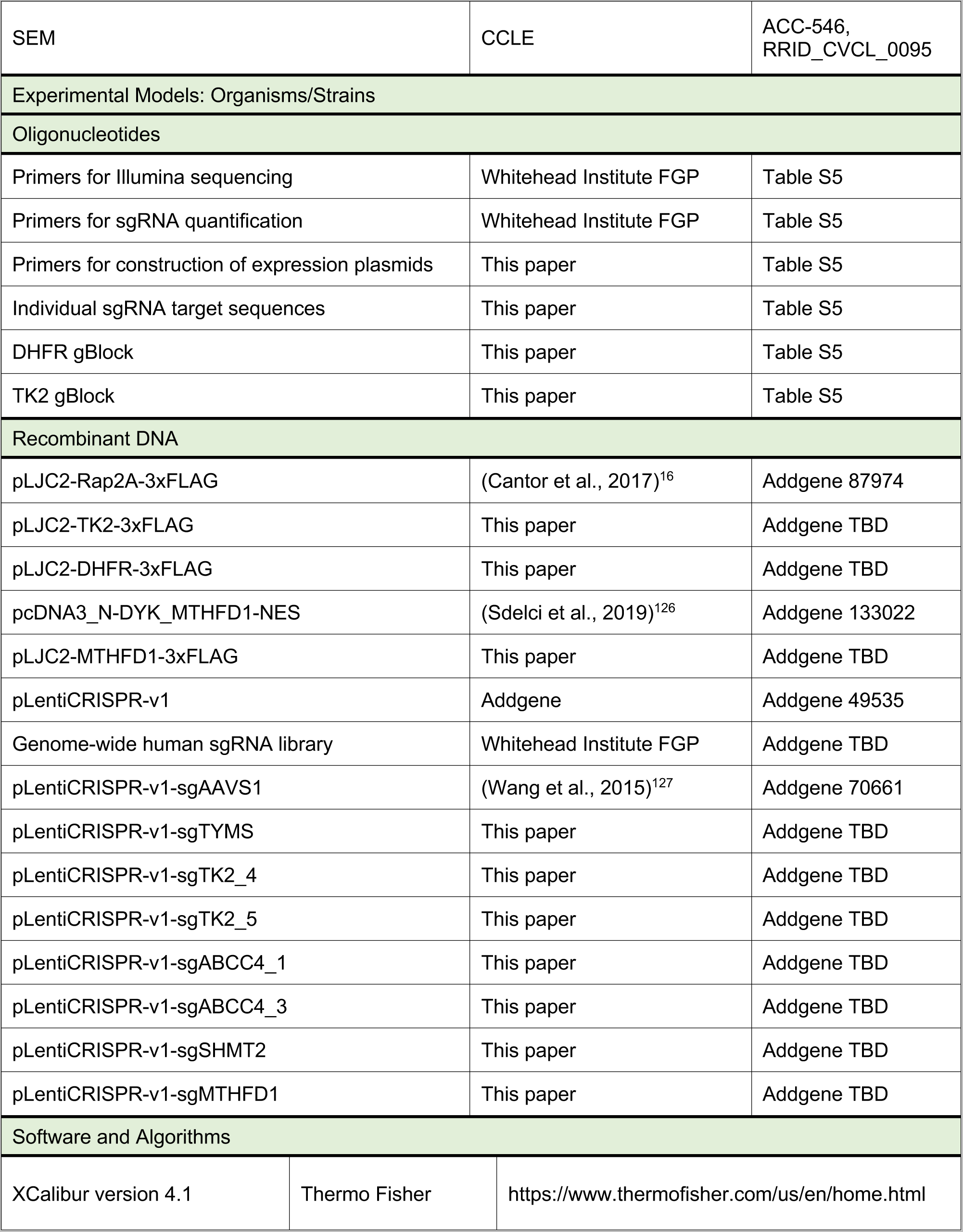
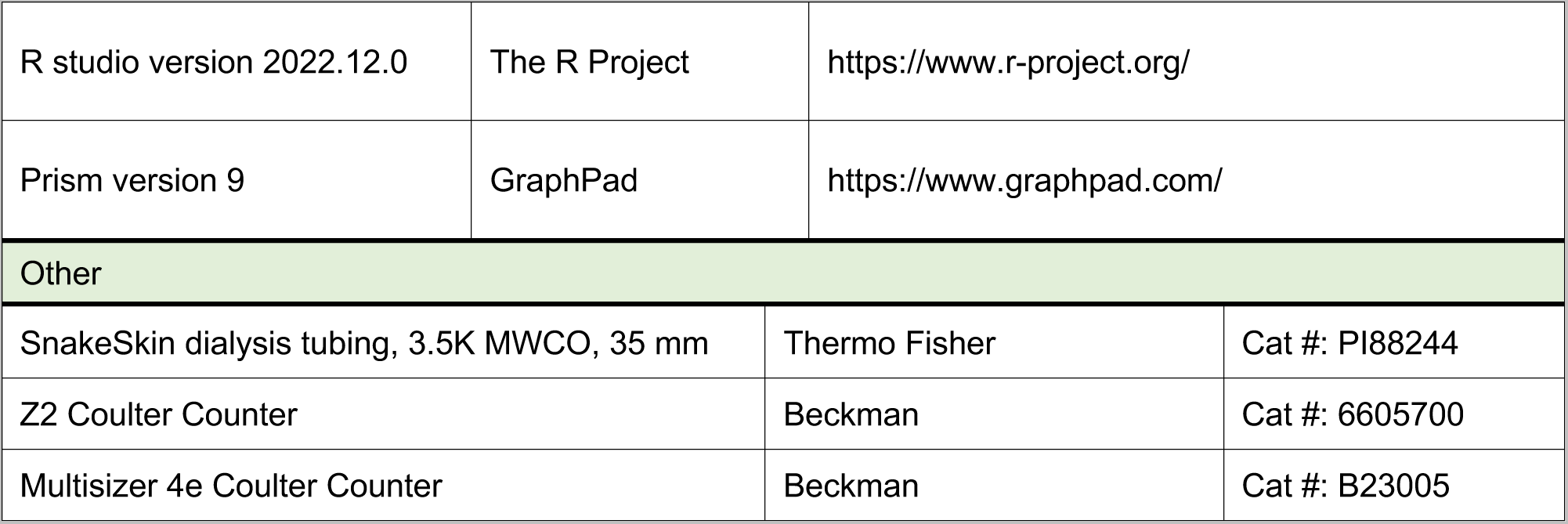

## RESOURCE AVAILABILITY

### Lead Contact

Further information and requests for resources and reagents should be directed to and will be fulfilled by the Lead Contact, Jason R. Cantor.

### Materials Availability

The individual gene knockout and expression plasmids generated in this study will be deposited in Addgene.

### Data and Code Availability

Datasets can be found in Tables S1, S3, and S4.

## EXPERIMENTAL MODEL AND SUBJECT DETAILS

### Cell lines

The following human cancer cell lines were kindly provided by: K562 and NOMO1, Dr. James Griffin (Dana Farber Cancer Institute); P12-Ichikawa, Dr. A. Thomas Look (Dana Farber Cancer Institute); and SEM, the Cancer Cell Line Encyclopedia (Broad Institute). Cell lines were verified to be free of mycoplasma contamination^128^ and their identities were authenticated by short tandem repeat (STR) profiling.

### Cell culture conditions

The following culture media were used in this study (all contained 0.5% penicillin-streptomycin):

1. RPMI^+S^: RPMI 1640, no glucose (Thermo Fisher) with 5 mM glucose and 10% FBS.
2. RPMI^+dS^: RPMI 1640, no glucose (Thermo Fisher) with 5 mM glucose and 10% dialyzed FBS.
3. RPMI11^+S^: RPMI 1640 (Thermo Fisher) with 10% FBS.
4. RPMI11^+2S^: RPMI 1640 (Thermo Fisher) with 20% FBS.
5. DMEM^+S^: DMEM, high glucose, GlutaMAX (Thermo Fisher) with 10% FBS.
6. DMEM^+2S^: DMEM, high glucose, GlutaMAX (Thermo Fisher) with 20% FBS.
7. HPLM^+dS^: homemade HPLM (See Table S2) with 10% dialyzed FBS and using RPMI 1640 100X Vitamins (Sigma-Aldrich R7256).
8. HPLM^+dS^: homemade HPLM (See Table S2) with 450 nM folic acid, eight other RPMI-defined vitamins, and 10% dialyzed FBS.
9. RPMI (manual)^+dS^: RPMI (See Table S2) with 5 mM glucose and 10% dialyzed FBS.

HPLM-based media using either (7) or (8) are distinguished with the shaded colors in Figures 3D and 3J. The basal HPLM formulation in (8) was used across all experiments except the chemical screens and those shown in Figures 3C, 3E, 3G, and S3B.

Using SnakeSkin tubing (Thermo Fisher PI88244), FBS was dialyzed as previously described^16^. Prior to use, all FBS-supplemented media were sterile filtered using a bottle-top vacuum filter with cellulose acetate membrane, pore size 0.2 μm (Corning 430626 or Nalgene 290-4520). Cultured cells were maintained at 37°C, atmospheric oxygen, and 5% CO_2_.

## METHOD DETAILS

### High-throughput chemical screens

NOMO1, P12-Ichikawa, and SEM cell lines were each initially maintained in RPMI^+S^. After four to six passages, cells were split and cultured in either RPMI^+S^, RPMI^+dS^, or HPLM^+dS^ for at least four passages prior to screening in each respective condition. Seeding densities for each cell line were tested and in turn selected to minimize growth rate differences between conditions. For each cell line-medium combination, cells were plated on the same day for high-throughput screening. Cells were seeded in 1536-well white solid bottom polystyrene Greiner plates containing 5 μL of growth medium using a Multidrop Combi Dispenser with a small tube dispensing cassette – the NOMO and SEM lines were plated at 500 cells/well and the P12-Ichikawa line at 1000 cells/well. Within each individual plate, one column was not seeded with cells and served as a background control. After seeding the cells, 23 nL of MIPE 4.1 library compounds (in DMSO) were added to each plate using a Kalypsys 1536 Pintool dispenser. Bortezomib (final concentration, 2 μM) and DMSO were similarly added to each plate as positive and negative controls for cytotoxicity, respectively. For each screen, MIPE 4.1 compounds were added over an eleven-point concentration range from 47 μM to 0.79 nM in 3-fold dilutions immediately after plating cells. Each plate was covered with a stainless steel gasketed lid to prevent evaporation and then housed in a tissue culture incubator maintained at 37°C, 95% relative humidity, and 5% CO_2_.

Following 48 hr incubation with compound, plate lids were removed and 3 μL of CellTiter-Glo One (Promega) were added to each well by using a solenoid valve dispenser. Plates were incubated with lids in place for 15 min at room temperature and then luminescence readings were taken using a ViewLux ultra HTS Microplate Imager (Perkin Elmer) with a 2 second exposure time per plate. Data for each compound were normalized as percent viability such that measurements for DMSO and the empty well controls in each plate were defined as 100% and 0%, respectively. Assay statistics were then calculated and all screened plates had a Z-factor greater than 0.4.

### Brivudine CRISPR modifier screens

#### Brivudine dose-responses in flask format

Following at least two passages in RPMI^+S^, K562 cells were pelleted and used to seed a T-75 cell culture flask (Corning 430641U) at a density of ∼150,000 cells/mL in 20 mL of HPLM^+dS^. After 48 hr incubation, pools of 2 million cells were used to seed T-75 culture flasks at a density of 100,000 cells/mL in 20 mL of HPLM^+dS^. Following 1 hr incubation of the seeded flasks, brivudine (BVDU) (final concentrations, 1 μM, 500 nM, 100 nM, or 50 nM) was added to the cells. All flasks, including the untreated controls, contained 0.25% DMSO. After 72 hr treatment, cell density measurements were recorded using a Coulter Counter (Beckman Z2 or Multisizer 4e) with a diameter setting of 8-30 μm and used to determine the EC_25_ for a culture flask format with relevance to the CRISPR screens. The stock solution of BVDU was prepared at 40 mM in DMSO.

#### Human sgRNA library amplification

pLentiCRISPRv2-Opti plasmid containing a human sgRNA library designed by Evgeni M. Frenkel was provided by the Whitehead Institute Functional Genomics Platform. The genome-wide human sgRNA library contained 98,077 constructs targeting 19,734 protein-coding genes and 208 non-coding RNAs (∼5 sgRNAs per target), and 499 total intergenic and non-targeting control sgRNAs. Library plasmid was transformed into *E. coli* Endura electrocompetent cells (Lucigen), plated onto prewarmed LB medium/agar containing 100 μg/mL carbenicillin in a 245 mm square bioassay dish (Corning 431111), and incubated for 16 hr at 30°C, yielding ∼10^8^ individual transformants – equivalent to ∼1,000-fold coverage of the theoretical library diversity. Colonies were scraped and pooled in LB medium, and plasmid DNA was extracted using an EndoFree Maxi Kit (QIAGEN).

#### Genome-wide CRISPR modifier screens

To achieve at least 1000-fold coverage of the sgRNA library following antibiotic selection, 300 million K562 cells were seeded at a density of 2.5 × 10^6^ cells/mL in 6-well plates containing 2 mL of RPMI11^+S^, 8 μg/mL polybrene, and the pLentiCRISPR-v2-Opti library virus. Spin infection was carried out by centrifugation at 2,200 RPM for 45 min at 37°C. After 18 hr incubation, cells were pelleted to remove virus and then re-seeded in fresh RPMI11^+S^ for 24 hr. Cells were then pelleted, re-seeded to a density of 150,000 cells/mL in RPMI11^+S^ containing 2 μg/mL puromycin (Sigma-Aldrich), and cultured for 72 hr. Following selection, an initial pool of 100 million cells was pelleted and frozen, and another pool of 175 million cells were used to collectively seed each of fourteen total 225 cm^2^ rectangular canted neck cell culture flasks (Corning 431082) at a density of 100,000 cells/mL in 125 mL of HPLM^+dS^. BVDU (final concentration, 100 nM) or DMSO was then added to each of seven flasks such that all cultures contained 0.25% DMSO. Cells were passaged every 72 hr, with fresh BVDU or DMSO added at each passage, and population doublings were tracked by cell density measurements using a Coulter Counter with a diameter setting of 8-30 μm. After 15 population doublings, a pool of 100 million cells from each screen was harvested for genomic DNA (gDNA) extraction using the QIAamp DNA Blood Maxi Kit (QIAGEN).

Using Ex Taq DNA Polymerase (Takara), sgRNA inserts from each initial and final pool were PCR-amplified from 144 μg of gDNA to achieve ∼400-fold coverage of the library. The PCR products were then purified and sequenced on a NextSeq 500 (Illumina) (Primer sequences are annotated in Table S5) to quantify sgRNA abundances in each sample.

### Plasmid construction

All oligonucleotides and gBlock Gene fragments used in this study are described in Table S5.

#### Construction of gene knockout plasmids

For each of the following genes, sense and antisense oligonucleotides were annealed and then cloned into *BsmBI*-digested pLentiCRISPR-v1: *TYMS, TK2, ABCC4, SHMT2,* and *MTHFD1*.

#### Construction of expression plasmids

The *TK2* and *DHFR* genes were amplified from codon-optimized gBlock gene Fragments (IDT) using the primers TK2-F/TK2-R and DHFR-F/DHFR-R, respectively, digested with *PacI*-*NotI*, and cloned into pLJC2-Rap2A-3xFLAG to generate pLJC2-TK2-3xFLAG and pLJC2-DHFR-3xFLAG. The *MTHFD1* gene was amplified from pcDNA3_N-DYK_MTHFD1-NES using the primers MTHFD1-F/MTHFD1-R, digested with *PacI*-*NotI*, and cloned into pLJC2-Rap2A-3xFLAG to generate pLJC2-MTHFD1-3xFLAG.

### Lentivirus production

To produce lentivirus, HEK293T cells in DMEM^+S^ were co-transfected with the VSV-G envelope plasmid, the Delta-VPR packaging plasmid, and transfer plasmid (either pLJC2, pLentiCRISPR-v1, or pLentiCRISPRv2-Opti backbone) using X-tremeGENE 9 Transfection Reagent (Sigma-Aldrich). The medium was exchanged with fresh DMEM^+2S^ 16 hr after transfection, and the virus-containing supernatant was collected at 48 hr post-transfection, passed through a 0.45 μm filter to eliminate cells, and then stored at −80°C.

### Cell line construction

#### Knockout cell lines

To establish *SHMT2*-knockout clonal cell lines, K562 cells were seeded at a density of 500,000 cells/mL in 6-well plates containing 2 mL RPMI11^+S^, 8 μg/mL polybrene, and pLentiCRISPR-v1 lentivirus. Spin infection was carried out by centrifugation at 2,200 RPM for 45 min at 37°C. After 16-18 hr incubation, cells were pelleted to remove virus and then re-seeded into fresh RPMI11^+S^ for 24 hr. Cells were then pelleted and re-seeded into RPMI11^+S^ containing puromycin for 72 hr. Following selection, the population was single-cell FACS-sorted into 96-well plates containing RPMI11^+2S^ (BD FACSMelody Cell Sorter). After 1.5-2 weeks, clones with the desired knockouts were identified by immunoblotting. To control for infection, a population of K562 cells was similarly selected following transduction with sgAAVS1-containing virus. The procedure to establish other knockout cell lines for short-term growth or drug treatment assays was similar except that cells were not FACS-sorted following puromycin selection (5 d post infection) (See **Short-term growth and drug treatment assays**).

#### TK2 cDNA expression cell line

To establish the *TK2* expression cell line, SEM cells were seeded at a density of 500,000 cells/mL in 6-well plates containing 2 mL of RPMI11^+S^, 8 μg/mL polybrene, and the pLJC2-TK2-3xFLAG lentivirus. Spin infection, culture medium exchange, and puromycin selection were carried out as described above for the knockout cell lines. Stable expression of *TK2* cDNA was confirmed by immunoblotting.

### Cell lysis for immunoblotting

Cells were centrifuged at 250 *g* for 5 min, resuspended in 1 mL ice-cold PBS, and then centrifuged again at 250 *g* for 5 min at 4°C. Cells were then immediately lysed with ice-cold lysis buffer (40 mM Tris-HCl pH 7.4, 1% Triton X-100, 100 mM NaCl, 5 mM MgCl_2_, 1 tablet of EDTA-free protease inhibitor (Roche 11580800; per 25 mL buffer), 1 tablet of PhosStop phosphatase inhibitor (Roche 04906845001; per 10 mL buffer). Cell lysates were cleared by centrifugation at 21130 *g* for 10 min at 4°C and quantified for protein concentration using an albumin standard (Thermo Fisher 23209) and Bradford reagent (Bio-Rad 5000006). Cell lysate samples were normalized for protein content, denatured upon the addition of 5X sample buffer (Thermo Fisher 39000), resolved by 12% SDS-PAGE, and transferred to a polyvinyl difluoride membrane (Millipore IPVH07850). Membranes were blocked with 5% nonfat dry milk in TBST for 1 hr at room temperature, and then incubated with primary antibodies in 5% nonfat dry milk in TBST overnight at 4°C. Primary antibodies to the following proteins were used at indicated dilutions: GAPDH (1:1000); RAPTOR (1:1000); TYMS (1:1000); SHMT2 (1:500); MTHFD1 (1:250); and TK2 (1:100).

Membranes were washed with TBST three times for 5 min each, and then incubated with species-specific HRP-conjugated secondary antibody (1:3000) in 5% nonfat dry milk for 1 hr at room temperature. Membranes were washed again with TBST three times for 5 min each, and then visualized with chemiluminescent substrate (Thermo Fisher) on a LICOR Odyssey FC.

### Cellular Thermal Shift Assay

Following at least one passage in HPLM^+dS^, 4 million K562 cells were pelleted and seeded to a density of 1 million cells/mL in 6-well plates containing 4 mL of HPLM^+dS^. After 5 min incubation of seeded plates, BVDU, fluorodeoxycytidine (FCdR), or methotrexate (MTX) (final concentration, 1 μM) was added to cells and then plates were gently shaken for 2 min. All wells, including the untreated controls, contained 0.25% DMSO. Following 2 hr treatment, cells were washed once with ice-cold PBS, pelleted, and then resuspended in 100 μL ice-cold PBS containing EDTA-free protease inhibitor (Roche). Cells were transferred to a PCR-strip tube and heated for 3 min using a Mastercycler Nexus X2 thermal cycler (Eppendorf). After heating, samples were snap-frozen with liquid nitrogen and then evenly thawed by transferring strip-tubes to a CoolRack XT (Corning) placed in a dry block heater held at 25°C. Following two additional freeze-thaw cycles with gentle pulse-vortexing after each thaw, cell lysates were cleared by centrifugation at 21130 *g* for 20 min at 4°C. Cell lysate samples were normalized for protein content and denatured upon the addition of 5X sample buffer.

The procedure used to determine the melting temperature for TYMS was similar to that described above with the following modifications:

1. Only K562 cells in HPLM^+dS^ with 0.25% DMSO were tested.
2. Cells were heated over a gradient of temperatures, either 37.1° to 49°C or 49.1° to 61.1°C, set using the Nexus X2.

### Short-term growth and drug treatment assays

#### Cell line panel, drug treatments

Following at least two passages in RPMI^+S^, cells were pelleted and used to seed T-25 cell culture flasks (Corning 430639) containing 12 mL of RPMI^+S^, RPMI^+dS^, or HPLM^+dS^. K562 cells were seeded to a density of ∼150,000 cells/mL, while the remaining cell lines (NOMO1, P12-Ichikawa, and SEM) were seeded at densities between 400,000-500,000 cells/mL. After 48 hr incubation, cells were pelleted and resuspended to densities of either 1 million cells/mL (K562, NOMO1, and P12-Ichikawa) or 2 million cells/mL (SEM) in the respective parent culture medium. From each resuspension, either 80,000 (K562, NOMO1, and P12-Ichikawa) or 160,000 (SEM) total cells were seeded in each of three replicate wells (per dosing concentration) in 6-well plates containing 4 mL of the appropriate culture medium. Following 1 hr incubation of seeded plates, compounds were added at specified doses and then plates were gently shaken for 2 min. All wells, including the untreated controls, contained 0.25% DMSO. After 96 hr treatment, cell density measurements were recorded using a Coulter Counter with a diameter setting of 8-30 μm. For each cell line compound combination, assays were performed across conditions in the same experiment.

#### Engineered K562 cell lines

The short-term growth and drug treatment assays with *SHMT2-*knockout K562 clonal cells were identical to those described above. For the *TYMS*-, *TK2-*, *ABCC4-*, and *MTHFD1*- knockout K562 cell lines, FACS-sorting was not performed following puromycin selection. Instead, 3 million cells were pelleted and resuspended to a density of 250,000 cells/mL in 12 mL of RPMI^+S^. After 48 hr incubation (7 d post infection), 3.5 million cells were pelleted and resuspended to a density of ∼300,000 cells/mL in 12 mL of HPLM^+dS^. Following 48 hr incubation (9 d post infection), pools of cells were pelleted and seeded for the 96 hr growth step – with drug treatment when appropriate – as described above, with cell density measurements ultimately recorded at 13 d post infection. To control for infection and all ensuing steps over the course of these two-week experiments, a population of K562 cells was transduced with sgAAVS1-containing virus, selected, passaged, and assayed parallel to unsorted genetic knockout populations.

The short-term treatment assay procedure using RPMI- or HPLM-based derivatives was identical to that above with minor modifications:

1. The following were added to the appropriate complete media only for the final 96 hr step of the assay: hypoxanthine, uridine, thymidine, cytidine, deoxyuridine, and deoxycytidine. Stock solutions of each individual component were prepared at 10 mM in either water or 0.2 M HCl (hypoxanthine and uridine).
2. Modified concentrations of amino acids, salt ions, and vitamins were incorporated at the preceding medium-specific 48 hr passage step. Stock solutions of individual vitamins were prepared as follows: biotin (8.2 mM in 20 mM NaOH), folic acid (450 μM in 20 mM NaOH), and 5-mTHF (1 mM in 50 mM NaOH).

#### K562 cells, for cellular folates

Following at least two passages in RPMI^+S^, K562 cells were pelleted and used to seed T-25 cell culture flasks at a density of ∼150,000 cells/mL in 12 mL of the appropriate HPLM-based medium. After 48 hr incubation, cells were pelleted and resuspended to a density 1 million cells/mL in the respective passage medium. From each resuspension, 500,000 total cells were seeded in each of three T-75 cell culture flasks containing 25 mL of the appropriate HPLM-based medium. After 1 hr incubation of seeded flasks, compounds were added at specified doses, and then treated for 96 hr. All flasks, including the untreated controls, contained 0.25% DMSO.

### Metabolite profiling and quantification of metabolite abundance

LC-MS analyses were performed on a QExactive HF benchtop orbitrap mass spectrometer equipped with an Ion Max API source and HESI II probe, coupled to a Vanquish Horizon UHPLC system (Thermo Fisher). External mass calibration was performed using positive and negative polarity standard calibration mixtures every 7 days. Acetonitrile was hypergrade for LC-MS (Millipore Sigma) and all other solvents were Optima LC-MS grade (Thermo Fisher).

#### Cells, polar metabolites

At the conclusion of short-term growth or treatment assays, a 500 μL aliquot from each well was used to measure cell number and volume via Coulter Counter with a diameter setting of 8-30 μm, and the remaining cells were centrifuged at 250 *g* for 5 min, resuspended in 1 mL ice-cold 0.9% sterile NaCl (Growcells MSDW1000), and again centrifuged at 250 *g* for 5 min at 4°C. Metabolites were extracted in 1 mL ice-cold 80% methanol containing 500 nM internal amino acid standards (Cambridge Isotope Laboratories). Following a 10 min vortex and centrifugation at 21130 *g* for 3 min at 4°C, samples were dried under nitrogen gas. Dried samples were stored at −80°C and resuspended in 100 μL water. After a 10 min vortex and centrifugation at 21130 *g* for 10 min at 4°C, 2.5 μL from each cell sample was injected onto a ZIC-pHILIC 2.1 x 150 mm analytical column equipped with a 2.1 x 20 mm guard column (both were 5 μm particle size, Millipore Sigma). Buffer A was 20 mM ammonium carbonate, 40 mM ammonium hydroxide; buffer B was acetonitrile. The chromatographic gradient was run at a flow rate of 0.15 mL/min as follows: 0-20 min: linear gradient from 80% to 20% B; 20-20.5 min: linear gradient from 20% to 80% B; 20.5-28 min: hold at 80% B. The mass spectrometer was operated in full scan, polarity-switching mode with the spray voltage set to 3.0 kV, the heated capillary held at 275°C, and the HESI probe held at 350°C. The sheath gas flow rate was set to 40 units, the auxiliary gas flow was set to 15 units, and the sweep gas flow was set to 1 unit. The MS data acquisition in positive mode was performed in a range of 50-750 m/z, with the resolution set to 120,000, the AGC target at 10^6^, and the maximum integration time at 20 msec. The settings in negative mode were the same except that the range was instead 70-1000 m/z.

For the highly targeted analysis of several metabolites, additional tSIM (targeted selected ion monitoring) scans were added with the following settings: resolution set to 120,000, an AGC target of 10^5^, maximum integration time of 200 msec, and isolation window of 1.0 m/z. The target masses, each in negative ionization mode, were: 87.0088 (pyruvate), 321.0493 (dTMP), 347.0398 (IMP), 480.982 (dTTP). For BVDU-treated samples, identical tSIM scans in negative ionization mode were added for the following target masses: 330.9935 (BVDU) and 410.9598 (BVDU-MP). For FCdR-treated samples, an identical tSIM scan in negative ionization mode was added for the following target mass: 325.0243 (FdUMP).

#### Media, relative folic acid availability

Samples of HPLM-based media containing either 450 nM or 2.27 μM folic acid were snap-frozen in liquid nitrogen and stored at −80°C prior to inoculation of short-term BVDU treatment assays. At the conclusion of these assays, a 500 μL aliquot from each well was collected and centrifuged at 250 *g* for 5 min. Metabolites were extracted from both the resulting supernatants and the initial samples by diluting 1:40 in a solution of 50:30:20 methanol:acetonitrile:water (MeOH:ACN:H_2_O) containing 500 nM internal amino acid standards. Following a 10 min vortex and centrifugation at 21130 *g* for 5 min at 4°C, 2 μL of each sample was injected for analysis as described above for profiling cell samples. For the highly targeted analysis of folic acid, an additional tSIM scan was added with the same settings described for cell samples, except that the scan was run in positive ionization mode and the target mass was 442.1470.

#### Comparison of vitamin abundances

To extract metabolites from basal RPMI (Thermo 11879), samples were diluted 1:10 into 50:30:20 MeOH:ACN:H_2_O containing 500 nM internal amino acid standards. For RPMI 1640 100X vitamins solution (Sigma R7256, Lots RNBB7627 and RNBK1269), aliquots were first diluted 1:10 in water and vortexed for 1 min at 4°C, and then metabolites were extracted by further diluting 1:10 into 50:30:20 MeOH:ACN:H_2_O containing 500 nM internal amino acid standards. Following a 10 min vortex and centrifugation at 21130 *g* for 5 min at 4°C, 2 μL of each sample was injected for analysis as described above for profiling cell samples. For the highly targeted analysis of biotin, an additional tSIM scan was added with the same settings described for cell samples, except that the scan was run in positive ionization mode and the target mass was 245.0954.

#### Serum

To extract metabolites from untreated and dialyzed FBS, samples were diluted 1:40 into 50:30:20 MeOH:ACN:H_2_O containing 500 nM internal amino acid standards. Following a 10 min vortex and centrifugation at 21130 *g* for 3 min at 4°C, samples were dried under nitrogen gas. Dried samples were stored at −80°C and resuspended in 100 μL water. After a 10 min vortex and centrifugation at 21130 *g* for 10 min at 4°C, 2 μL of each sample was injected for analysis as described above for profiling cell samples. For the highly targeted analysis of pyruvate, an additional tSIM scan was added with the same settings described for cell samples.

#### Cells, folates

At the conclusion of short-term treatment assays, a 500 μL aliquot from each flask was used to measure cell number and volume via Coulter Counter with a diameter setting of 8-30 μm. From each flask, 3 million cells were then centrifuged at 250 *g* for 5 min, resuspended in 1 mL ice-cold 0.9% sterile NaCl, and again centrifuged at 250 *g* for 5 min at 4°C. The ensuing protocol for folate extraction was adapted from others described elsewhere^95, 129, 130^. Metabolites were extracted in 1 mL ice-cold 80% methanol containing 2.5 mM sodium ascorbate, 25 mM ammonium acetate pH 7, and 100 μm aminopterin. Following a 10 min vortex and centrifugation at 21130 *g* for 10 min at 4°C, samples were dried under nitrogen gas. Dried samples were stored at −80°C and reconstituted in 400 μL ice-cold resuspension buffer (50 mM K_2_HPO_4_ pH 7, 30 mM ascorbic acid, 0.5% 2-mercaptoethanol). Following a 10 min vortex and centrifugation at 21130 *g* for 10 min at 4°C, resulting supernatants were transferred to fresh tubes with 25 μL charcoal-treated rat serum, and then gently shaken at 300 RPM for 2 hr at 37°C using a Thermomixer C (Eppendorf). Sample pH was then adjusted to 4 with 15 μL of 20% formic acid before loading onto conditioned SPE Bond-Elute pH columns (Agilent, 14102062) at 4°C. After washing with 1 mL aqueous buffer (25 mM ammonium acetate pH 4, 30 mM ascorbic acid), samples were eluted in 400 μL 50% methanol containing 30 mM ammonium acetate pH 7 and 0.5% 2-mercaptoethanol, and then dried under nitrogen gas. Dried samples were resuspended in 50 μL water. After a 10 min vortex and centrifugation at 21130 *g* for 10 min at 4°C, 5 μL of each sample was injected for analysis as described above for profiling cell samples. For the highly targeted analysis of folate species, additional tSIM scans were added with the same settings described for cell samples, except that the scans were run in positive ionization mode and the target masses were: 446.1783 (THF), 460.1939 (5-mTHF), and 474.1732 (10-formyl-THF). Peaks corresponding to 5-formyl-THF and 10-formyl-THF were distinguished based on retention time (See Table S3).

To generate charcoal-treated rat serum for cleaving polyglutamate tails from intracellular folates, 250 mg activated charcoal (Sigma-Aldrich C9157) was added to 5 mL rat serum (Sigma-Aldrich R9759) and then incubated head-over-head for 3 hr at 4°C. After centrifugation at 1500 *g* for 10 min at 4°C, supernatants were collected and stored at −20°C. To condition SPE Bond-Elute columns, 1 mL LC-MS grade methanol and 1 mL aqueous buffer were sequentially loaded.

#### Enzyme activity assay evaluation

For the detection of metabolites from DHFR, MTHFD1-DC, MTHFD1-S, and trifunctional MTHFD1 activity assays, reaction mixtures were extracted (See **Enzyme activity assays**) and 5 μL of each sample was injected for analysis as described above for profiling cell samples but using the following chromatographic gradient: 0-10 min: linear gradient from 80% to 20% B; 10-10.5 min: linear gradient from 20% to 80% B; 10.5-17.5 min: hold at 80% B. For the highly targeted analysis of reaction components in activity assays, additional tSIM scans were respectively added with the same settings described for cell samples, except that the target masses were: 426.0221 (ADP), 474.1732 (10-formyl-THF, positive ionization mode), and 742.0682 (NADP).

#### Synthesis of brivudine monophosphate

For isolation of BVDU-MP from in vitro TK2 reactions, the twice-dried samples were resuspended in 100 μL water (See **Enzyme activity assays**). Following a 10 min vortex and centrifugation at 21130 *g* for 3 min at 4°C, 5 μL aliquots were injected onto the LC-MS with the same settings and chromatographic gradient described above for profiling cell samples. Based on an empirically determined retention time of ∼6.6 min for BVDU-MP, fractions from eight successive sample injections were collected from 5-8 min by disconnecting the viper fitting that feeds into the MS. Fractions were dried under nitrogen gas and stored at −80°C. Dried samples were reconstituted in 100 μL water. Following a 10 min vortex and centrifugation at 21130 *g* for 3 min at 4°C, the supernatants were pooled and then dried under nitrogen gas. The dried sample was resuspended in 100 μL water and stored at −80°C. Purified BVDU-MP and a 100 μM stock solution of NMP chemical standards (AMP, CMP, GMP, IMP, dTMP, and UMP) were diluted 1:10 in water containing internal amino acid standards. Following a 10 min vortex and centrifugation at 21130 *g* for 3 min at 4°C, 5 μL of each sample was injected for analysis. The concentration of BVDU-MP was estimated as the average of normalized peak areas across the NMPs.

#### Identification and quantification

Metabolite identification and quantification were performed with XCalibur version 4.1 (Thermo Fisher) using a 10-ppm mass accuracy window and 0.5 min retention time window. To confirm metabolite identities and to enable quantification when desired, a manually constructed library of chemical standards was used. Standards were validated by LC-MS to confirm that they generated robust peaks at the expected m/z ratio, and stock solutions were stored in pooled format at −80°C. On the day of a given queue, stock solutions were diluted 1:10 in either water (cell samples) or 50:30:20 MeOH:ACN:H_2_O (remaining samples) containing 500 nM internal amino acid standards, and then vortexed and centrifuged as described for biological samples. Aminopterin served as the internal standard for profiling cellular folates. For those metabolites lacking a standard, peak identification was restricted to high confidence peak assignments. See Table S3.

Given that metabolite extraction protocols differed by sample type, the internal standard concentrations in processed samples for polar metabolites varied: chemical standards (450 nM), basal RPMI and RPMI 100X vitamins solution (450 nM), media and serum samples (487.5 nM), and cell samples (5 μM). Therefore, the raw peak areas of internal standards within each sample of a given batch were first normalized to account for these differences. Metabolite quantification was performed as described elsewhere. For the final concentrations of chemical standards used to quantitate specific metabolites, see Table S3.

### Synthesis of 10-formyl-tetrahydrofolate

To synthesize 10-formyl-THF, a reported protocol was adapted^131^. Reactions containing 20 mM 5,10-me^+^-THF (Schircks 16.230) and 70 mM 2-mercaptoethanol were carried out in assay buffer (100 mM Tris-HCl pH 8.5) in a total volume of 100 μL. After a 1 min vortex at 37°C, reactions were incubated for 1 hr at 25°C. Assuming complete substrate conversion, aliquots of 20 mM 10-formyl-THF were then stored at −80°C. The stock solution of 5,10-me^+^-THF was prepared at 100 mM in DMSO. Tris-HCl buffer was prepared in LC-MS grade water and the pH adjusted using KOH.

### Expression and immunoprecipitation of recombinant proteins

For isolation of recombinant proteins, 4 million HEK293T cells were plated in 15 cm culture dishes containing DMEM^+S^. After 24 hr incubation, cells were transfected with 15 μg of pLJC2 constructs harboring TK2-3xFLAG, DHFR-3xFLAG, or MTHFD1-3xFLAG as described elsewhere. Following an additional 48 hr incubation, cells were rinsed once with ice-cold PBS and then immediately lysed in ice-cold lysis buffer (See **Cell lysis for immunoblotting**). Cell lysates were cleared by centrifugation at 21130 *g* for 10 min at 4°C. For anti-FLAG immunoprecipitation, FLAG-M2 affinity gel (Sigma-Aldrich) was washed three times in lysis buffer, and then 400 μL of a 50:50 affinity gel slurry was added to a pool of clarified lysates collected from either five (MTHFD1-3xFLAG) or ten (TK2-3xFLAG and DHFR-3xFLAG) individual 15 cm culture dishes, and incubated with rotation for 3 hr at 4°C. Following immunoprecipitation, the beads were washed twice in lysis buffer and then four times with lysis buffer containing 500 mM NaCl. Recombinant protein was then eluted in lysis buffer containing 500 μg/mL 3x-FLAG peptide (Sigma-Aldrich) for 1 hr with rotation at 4°C. The eluent was isolated by centrifugation at 100 *g* for 4 min at 4°C (Bio-Rad 732-6204), buffer exchanged (Amicon Ultra 10 kDa MWCO UFC501024, TK2-3xFLAG and DHFR-3xFLAG; 30 kDa MWCO UFC503024, MTHFD1-3xFLAG) against 20 volumes of storage buffer (40 mM Tris-HCl pH 7.5, 100 mM NaCl, 2 mM dTT), mixed with glycerol (final concentration 15% v/v), snap-frozen with liquid nitrogen, and stored at −80°C.

Protein samples were quantified using an albumin standard and Bradford reagent. Purified proteins were denatured upon the addition of 5X sample buffer and resolved by 12% SDS-PAGE.

### Enzyme activity assays

For all enzyme reactions, the assay buffer was 40 mM Tris-HCl pH 7.4, 5 mM Na_2_HPO_4_, 5 mM MgCl_2_, 2 mM dTT, 100 μM NaCl.

#### TK2, for synthesis of brivudine monophosphate

To synthesize BVDU-MP, ten parallel reactions of purified recombinant TK2 (10-20 nM) with ATP (2 mM) and BVDU (2 mM) in assay buffer were carried out in PCR strip-tubes (100 μL) for 12 hr at 37°C using a Nexus X2 thermal cycler. After pooling the reactions, metabolites were extracted by transferring 330 μL aliquots to fresh tubes containing 770 μL 50:30:20 MeOH:ACN:H_2_O. After a 10 min vortex and centrifugation at 21130 *g* for 3 min at 4°C, supernatants were transferred to fresh tubes and dried under nitrogen gas. Dried samples were resuspended in 125 μL water and, after a 10 min vortex and centrifugation at 21130 *g* for 3 min at 4°C, supernatants were pooled and dried under nitrogen gas. For isolation of purified BVDU-MP, see **Metabolite profiling and quantification of metabolite abundance**.

#### DHFR

Reactions of recombinant DHFR (20-40 nM) with NADPH (100 μM), DHF (100 μM), and either MTX or BVDU-MP (10 μM) in assay buffer were carried out at 37°C in a total volume of 50 μL. Following 5 min incubation, a 30 μL aliquot of the reaction was removed and immediately added to 70 μL ice-cold MeOH:ACN:H_2_O containing 500 nM internal amino acid standards for metabolite extraction. Samples were then vortexed for 5 min and centrifuged at 21130 *g* for 1 min at 4°C.

NADP levels generated in each reaction were evaluated by LC-MS analysis of extracted samples. An identically prepared extraction sample containing only the two DHFR substrates was used to correct for background NADP. Using a NADP chemical standard (10 μM), we determined that reactions without MTX achieved ∼40-45% turnover.

#### MTHFD1-DC

Reactions of recombinant MTHFD1 (40-60 nM) with NADPH (100 μM), 10-formyl-THF (100 μM), and either LY-345899 or BVDU-MP (10 μM) in assay buffer were carried out at 37°C in a total volume of 50 μL. Following 15 min incubation, a 30 μL aliquot of the reaction was removed and immediately added to 70 μL ice-cold MeOH:ACN:H_2_O containing 500 nM internal amino acid standards for metabolite extraction. After a 5 min vortex, samples were centrifuged at 21130 *g* for 1 min at 4°C.

NADPH, NADP, and 10-formyl-THF levels from each reaction were evaluated by LC-MS analysis of extracted samples. An identically prepared sample containing only the MTHFD1-DC substrates was used to correct for background NADP. Using a NADP chemical standard (10 μM), we determined that reaction samples without LY-345899 achieved ∼35-40% turnover.

#### MTHFD1-S

Reactions of purified recombinant MTHFD1 (40-60 nM) with ATP (100 μM), THF (100 μM), and formate (1 mM) in assay buffer were carried out at 37°C in a total volume of 50 μL. Following 15 min incubation, a 30 μL aliquot of the reaction was removed and immediately added to 70 μL ice-cold MeOH:ACN:H_2_O containing 500 nM internal amino acid standards for metabolite extraction. After a 5 min vortex, samples were centrifuged at 21130 *g* for 1 min at 4°C. ADP and 10-formyl-THF levels in each reaction were evaluated by LC-MS analysis of extracted samples. However, by evaluating an identically prepared sample containing only the MTHFD1-S substrates to correct for background ADP and 10-formyl-THF, we found that synthesis of each product was negligible.

#### Trifunctional MTHFD1

Reactions of recombinant MTHFD1 (40-60 nM) with ATP (100 μM), THF (100 μM), formate (1 mM), NADPH (100 μM), and a non-endogenous compound (10 μM; LY-345899, BVDU-MP, BVDU, or MTX) in assay buffer were carried out at 37°C in a total volume of 50 μL. Following 15 min incubation, a 30 μL aliquot of the reaction was removed and immediately added to 70 μL ice-cold MeOH:ACN:H_2_O containing 500 nM internal amino acid standards for metabolite extraction. After a 5 min vortex, samples were centrifuged at 21130 *g* for 1 min at 4°C.

Abundances of the following reaction components were evaluated by LC-MS analysis of extracted samples: ATP, ADP, 10-formyl-THF, NADPH, and NADP. An identically prepared sample containing only the four substrates was used to correct for background ADP and NADP.

Stock solutions of compounds used in enzyme activity assays were prepared as follows: ATP (20 mM in water), THF (100 mM in water), ammonium formate (100 mM in water), NADPH (20 mM in 10 mM NaOH), DHF (20 mM in 500 mM NaOH), MTX (40 mM in DMSO), LY-345899 (40 mM in DMSO), and BVDU (40 mM in DMSO).

## QUANTIFICATION AND STATISTICAL ANALYSIS

### High-throughput chemical screens

Normalized viability data from the NCATS screening platform were fit to a 4-parameter log-logistic model using the *drc* package in R to generate dose-response curves^132^. For two compounds duplicated in the MIPE 4.1 library (NCGC00160391 and NCGC00179501), viability values at each concentration were averaged prior to similarly fitting the data, resulting in 17,784 cell line-medium-compound combinations. A subset of 654 curves (3.6% of the total dataset) failed to converge when fit to the logistic model. These were largely associated with compounds that had minimal effects on viability but also showed high data variability that prevented model convergence. To obtain metrics for further downstream analysis, this subset was instead fit using linear regression.

Areas under the dose-response curve (AUC) were calculated using the trapezoidal rule at the eleven log_10_-transformed dosing concentrations. Most fitted curves showed maximum values greater than the untreated controls used for normalization. Therefore, to reduce potential false positives in calculating differential AUC values between screens, curves with a maximum viability greater than 100% – and the corresponding curve metrics (AUC, minimum viability, and residual standard error) – were scaled by the maximum curve value. Moreover, a small subset of curves also exhibited a sharp decrease in viability over the two lowest dosing concentrations, in turn likely generating artifacts with this scaling method. Therefore, a subset of 73 curves that exhibited a maximum value greater than 100% and a 30% decrease in viability over the two lowest doses were refit to a linear model as well.

Residual standard error (RSE) distributions varied by cell line, with the SEM line exhibiting the largest median RSE. Thus, to again minimize potential false positives, curves with RSE values that were above cell line-dependent 98^th^ percentiles following viability scaling were removed from further downstream analysis. Next, to minimize potential false positives due to variance in initial viability measurements rather than to compound activity, cell line-specific curves with greater than 15% differences in maximum viability between two or more conditions for a given compound were also removed from further downstream analysis. At this point, filtered compounds not represented over all three conditions across cell lines were removed as well. Our collective curve fitting and filtering strategies established cell line-specific sets of compounds remaining for all three media: NOMO1 (1,871), SEM (1,761), and P12-Ichikawa (1,894). From these cell line-specific sets, 1,638 total compounds were common across cell line-medium combinations. Lastly, a set of 500 pan-inactive compounds were defined on the basis of exhibiting minimum scaled values greater than respective median scaled values in each of the nine screen datasets.

Response scores for the 1,138 shared active compounds were defined as corresponding AUCs. For each compound, differential response scores in each cell line were calculated between each pairwise set of conditions and then standardized (Z-score) relative to the entire set of 1,138 active compounds to assess differential sensitivity. For each compound, the differential sensitivity scores were then averaged across cell lines for each pairwise set of conditions.

### Genome-wide CRISPR screens

CRISPR screen analysis was performed as previously described^30, 133^. Sequencing reads were aligned to the sgRNA library to generate read counts and only exact matches were allowed. sgRNAs with less than 50 counts in the initial population were removed from further downstream analysis. Genes targeted by less than four distinct sgRNAs following this filtering process were also removed. The relative abundances of all remaining sgRNAs were determined by adding a pseudocount of one and then normalizing to the total reads in the sample. Depletion scores were calculated as the log_2_ fold-change in sgRNA abundance between the initial population and each final population. Gene scores were defined as the average log_2_ fold-change in depletion scores of all sgRNAs targeting the gene.

Screens performed in different conditions may introduce discrepancies in aggregate gene selection that affect the dynamic range of gene scores^134^. Therefore, to reduce potential bias in calculating differential scores based on assuming that such distributions are equivalent between screens, we scaled all gene scores instead based on the assumption that the sets of nontargeting (NT) sgRNAs and core essential genes (CEGs) would exhibit the same selection across different screens. Gene scores were scaled such that the medians of post-filtering NT sgRNAs (449) and reference CEGs (682 genes)^79^ included in the library were defined as 0 and −1, respectively, using the following equation where *X_S_* is the scaled gene score:

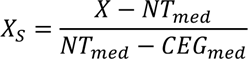

For each gene, a differential score was calculated between the two screening conditions and then standardized (Z-score) relative to the entire set of differential scores.

### Probability of dependency for genome-wide CRISPR screens

For each genome-wide screen, probabilities of dependency (PODs) were calculated for all library targets^135^. In brief, the gene score dataset from each screen was treated as a mixture model comprised of two normal distributions – distinct sets of non-essential and essential genes, with the latter having the lower mean. Densities were generated using a standard E-M optimization procedure initialized with parameters (mean, standard deviation, proportional value) of (−1, 0.3, 0.1) and (−0.2, 0.15, 0.9) for the reference sets of essential and non-essential genes, respectively. These initial values were based on empirical observations of score distributions for CEGs and nonessential genes from previous screens^30, 79^. The POD for a given gene was then calculated as the ratio of CEG density to the sum of the two densities at the gene score of interest. Given that standard deviations of the two distributions differ, their estimated densities converge to zero at different rates in tail regions, which can cause erroneous inflation of estimated probabilities at large enough gene score values. Thus, we identified the minimum POD and its corresponding gene score in each screen, and in turn, assigned the minimum probability to all targets with a gene score greater than that value.

### Receiver-operator analysis

For each CRISPR screen dataset, receiver-operator characteristic (ROC) curves were generated from relatively balanced reference sets of 682 CEG and 879 nonessential genes^79^. Area under the ROC curve was used as the performance metric to assess how well gene scores in each dataset could discriminate for CEGs.

*P*-values to compare cell density measurements and relative metabolite levels were determined using a two-tailed Welch’s *t*-test. The exact value of *n* and the definition of center and precision measures are provided in associated figure legends. Bar graphs were prepared in GraphPad Prism 9; remaining plots and heatmaps were prepared in R. All instances of reported replicates refer to *n* biological replicates.

